# A glucocorticoid-responsive polygenic signature in the anterior cingulate cortex moderates the association of early-life adversity and vulnerability for depression

**DOI:** 10.64898/2026.01.14.699350

**Authors:** Danusa Mar Arcego, Irina Pokhvisneva, Jan-Paul Buschdorf, Nicholas O’Toole, Sachin Patel, Randriely Merscher Sobreira de Lima, Guillaume Elgbeili, Patrick Lee, Gabriella Frosi, Laura Fiori, Corina Nagy, Patrícia Pelufo Silveira, Gustavo Turecki, Michael J. Meaney

## Abstract

Stress exposure is a major risk factor for psychopathology, yet how stress mediators shape long-term psychiatric vulnerability in humans remains unclear. Glucocorticoids, central effectors of the stress response, regulate gene expression through tissue-specific transcriptional programs, suggesting that glucocorticoid-responsive networks may shape sensitivity to adversity. Using RNA-sequencing following chronic glucocorticoid exposure in a non-human primate model, we identified a gene co-expression network specific to the anterior cingulate cortex (ACC) that was highly preserved across human post-mortem brain datasets relevant to depression. We derived an expression-based polygenic score (ePGS) reflecting genetic variation in network activity and tested its interaction with adversity in the UK Biobank. The ACC-specific glucocorticoid-responsive ePGS moderated the association between adversity and depressive symptoms in adult females, with the strongest effects for early-life adversity. Network genes were enriched for neurodevelopmental processes and showed stronger co-expression during childhood, highlighting a developmentally sensitive, region-specific mechanism linking stress exposure to depression risk.

## INTRODUCTION

Stressors are major triggers for psychopathology. The primary mediators of the stress response signals are the catecholamines and glucocorticoids (GCs) that promote adaptation through actions at a wide range of tissues ^1–5^. There is compelling evidence for GC effects on mental health. GCs are widely-prescribed treatments, particularly for auto-immune conditions ^6^.

However, prolonged GC treatment significantly increases the risk for mental disorders ^7,8^ including depression, anxiety and suicide ^9,10^. The risk is dose-related and commonly resolved with discontinuation ^7^. Likewise, the hypercortisolemia accompanying Cushing’s Disease bears a dramatic risk for depression, anxiety, mania and psychosis ^11,12^. Cushing’s Disease alters neural structure, including decreased gray matter volumes in the hippocampus and anterior cingulate cortex, as well as alterations in white matter in corticolimbic regions ^13^. The psychiatric conditions and structural alterations correlate with the severity of hypercortisolemia and are at least partially reversed with successful surgical intervention ^14,15^.

Genomic analyses corroborate the role of GC-signaling in mental disorders ^16^. A large post-mortem, multiomic analysis of human prefrontal cortex (PFC), hippocampus and amygdala ^17^ identified GC-responsive pathways within neurons and glia in individuals with major depressive disorder (MDD), with the strongest finding in the PFC. The glucocorticoid receptor (GR) emerges from such analyses as a prominent transcriptional regulator across the various omics data sets for MDD. Genetic variants that moderate the regulatory impact of GR are enriched for variants associated with psychiatric disorders ^18–20^. Omics analyses further implicate the GR co-chaperone, *FKBP5* ^21^ that regulates intra-cellular GR signaling. *FKBP5* exhibited differential expression in PFC excitatory and inhibitory neurons and oligodendrocytes for MDD ^17,22^, consistent with findings from snRNA-seq from MDD samples in the PFC of increased *FKBP5* expression in excitatory neurons ^23^.

Evidence for the importance of GC signaling thus emerges from diverse research models. GCs signal through the GR, a ligand-gated transcriptional factor, to regulate thousands of genes with precise tissue-specificity ^24^. Transcriptional analysis across 8 corticolimbic mouse brain regions following treatment with dexamethasone, a potent GR agonist, revealed that less than 10% of the GC-induced differentially-expressed genes (DEGs) were shared across regions and almost 50% were unique to a single region ^25^. An understanding of the effects of stress mediators on mental health, as well as the ability to leverage this knowledge towards novel therapeutics, will therefore require analyses of the highly tissue-specific GC signaling pathways.

As noted above, genomic analyses underscore the relevance of GC-signaling in the PFC, including the anterior cingulate cortex (ACC). GC signaling through the GR is well-established in the PFC ^26,27^. Neuroimaging studies consistently identify changes in activity within the ACC associated with MDD ^28–33^ and that are reversed with successful antidepressant treatment ^34–36^ including that of ketamine ^37^. The human neuroimaging findings together with those from clinical and genomic analyses noted above suggest that GC-induced effects on gene expression in the ACC may moderate the effects of stressors on the risk for depression. The challenge is that of devising approaches to test such associations in humans that allows a focus on tissue-specific signaling. MDD is highly polygenic ^38^ and thus associated with changes in the expression of multiple genes that operate within networks in the regulation of biological pathways to influence relevant neural functions. A gene network approach is thus best positioned to inform on relevant molecular mechanisms and biological processes ^39,40^.

The established role of GCs as mediators of the effects of stressors suggested that GR-regulated gene networks might moderate the effects of adversity on vulnerability for psychopathology ^41^. We reasoned that the transcriptomic-based identification of an ACC-specific, GC-regulated co-expression gene networks could be marshalled to examine the role of GC-signaling in moderating the effects of adversity on the risk for MDD. Transcriptomic data, unlike that of genotype, allows for the identification of tissue-specific gene networks associated with targeted signals. Our approach involved RNA-sequencing (RNAseq) following chronic GC exposure to identify GC-sensitive gene networks in the ACC using a non-human primate model. The transcriptomic data were analyzed using Weighted Gene Co-expression Network Analysis (WGCNA)^42^ to identify ACC-specific, GC-responsive gene co-expression networks. One such ACC-specific network was both GC-responsive and highly preserved across multiple human post-mortem RNAseq data sets of relevance for MDD.

The challenge was to then adapt this ACC-specific, GC-responsive gene network derived from non-human primate RNAseq for analyses of human data sets. To do so, we first identified human orthologs of the genes in the non-human primate ACC GC-regulated gene network. We then calculated an expression-based polygenic score (ePGS) ^43–47^ which is comprised of single-nucleotide polymorphisms (SNPs) that increase the expression of the genes in the target gene network, specifically in the cingulate cortex (i.e., cis quantitative expression trait loci; eQTL’s). The rationale for this approach assumes that a cumulative index of functional SNPs associated with increased expression in genes in the ACC-specific GC-responsive network should reflect the capacity of the functional activity of the network in response to adversity-enhanced GC exposure. A higher network-specific ePGS should thus predict mental health outcomes under conditions of adversity. We then used data from the UK Biobank to show that, indeed, an ePGS generated from the polymorphisms in genes that comprised an ACC-specific, GC-sensitive gene network significantly moderated the association between adversity and the risk for MDD. Subsequent analyses revealed that the moderation of the effect of the ACC-specific GC-responsive gene network on the risk for MDD was 1) most notable in analyses of early life adversity, 2) that the network genes were highly enriched for biological processes associated with neurodevelopment and 3) that the co-expression of network genes was significantly greater in childhood than in later life.

## RESULTS

### Glucocorticoid-responsive genes

We characterized the transcriptional sensitivity to GCs focusing on gene expression changes in the ACC of adult female macaques following chronic exposure to betamethasone, a high-affinity GR agonist, or saline (**Figure 1A**). The animals for these studies derived from the closure of a breeding facility for which only adult females were available. RNA-seq analysis revealed 246 differentially expressed genes (DEGs) following betamethasone exposure, with 143 up- and 103 down-regulated genes (FDR<0.05, **Figure 1B**, and **Table S1**). The DEGs included well-established GC-responsive genes, including FKBP5, HSD11B1, and KLF9, all canonical targets of GR signaling. Additional relevant genes included IGF-related genes (IGFBP5 and IGFBP6), which participate in insulin-signaling pathways, consistent with the established interaction between GC exposure, metabolic function, and insulin sensitivity ^48,49^. Several downregulated genes involved in inflammation or immune activity were also identified, including *CD74*, *IFI44*, *C1QTNF3*, *IDO1*, and *UBD*, reflecting the classical immunosuppressive GC effects (**Figure 1C**).

**Figure 1.**
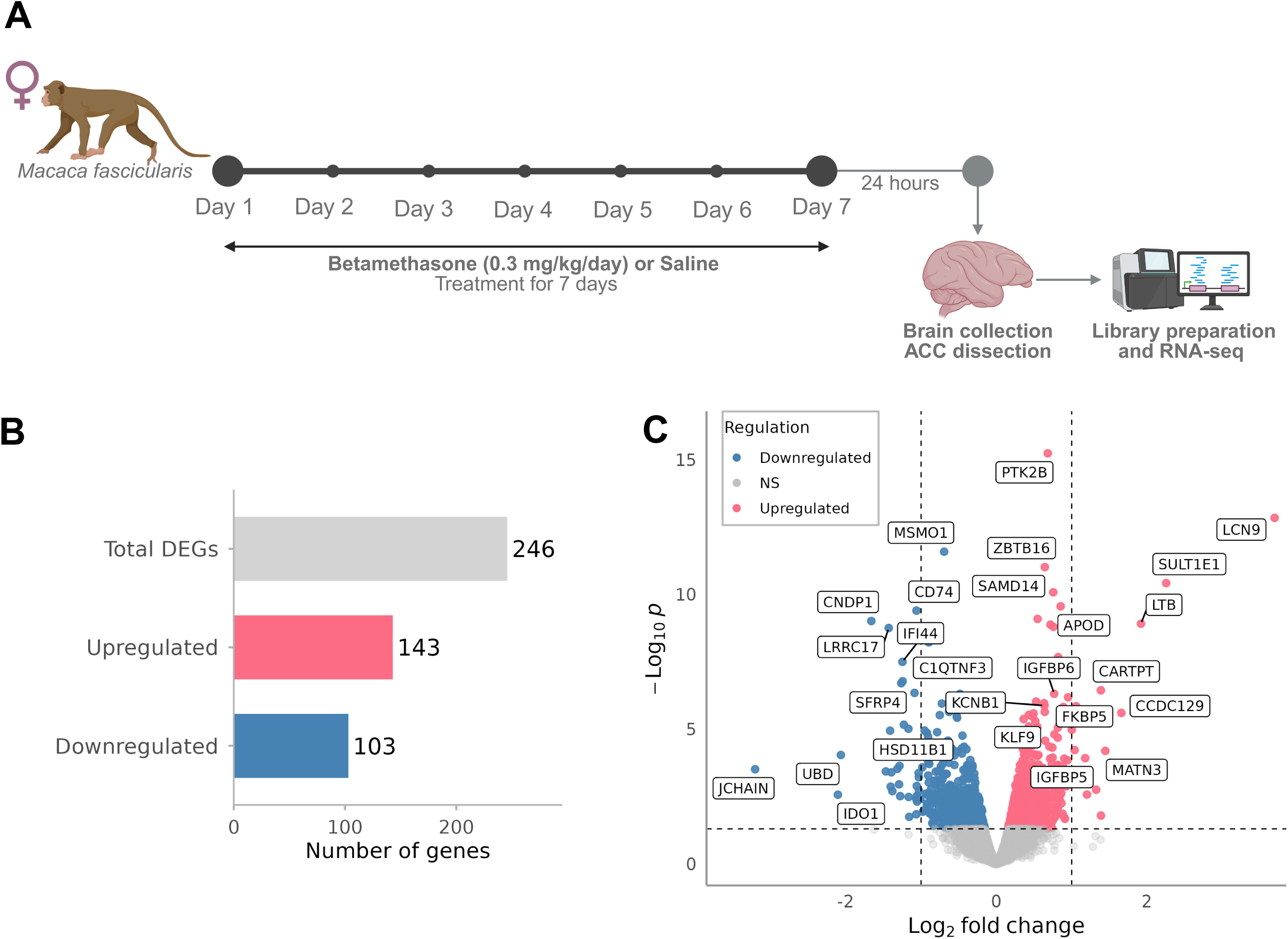
RNA-sequencing analysis workflow and differential gene expression in the macaque anterior cingulate cortex. **(A)** Experimental design and timeline of chronic betamethasone or saline treatment in female *Macaca fascicularis*, followed by anterior cingulate cortex (ACC) tissue collection and RNA-sequencing (RNA-seq). **(B)** Overview of differentially expressed genes (DEGs) identified in the ACC. **(C)** Volcano plot showing gene-level differential expression results. Upregulated genes are shown in pink, downregulated genes in blue, and non-significant genes in grey. Dashed lines indicate the statistical significance threshold and log_2_ fold-change cut-off.

Pathway enrichment analysis of DEGs (MetaCore, Clarivate Analytics) revealed that up-regulated genes were primarily enriched for pathways related to estradiol, adenosine, and adrenergic receptor signaling, oxidative stress responses and calcium-mediated signaling (**Figure 2A**). These pathways intersect with GC-mediated modulation of neuronal excitability, stress responses, and receptor sensitivity. Down-regulated genes were predominantly associated with immune processes and cholesterol biosynthesis (**Figure 2B**). Consistent with these patterns, gene ontology processes indicated that up-regulated genes were enriched for neuronal and synaptic functions, including regulation of synaptic signaling and neurotransmission (**Figure 2C**), whereas down-regulated genes associated with immune-related pathways and lipid and metabolic functions (**Figure 2D**). Gene Set Enrichment Analysis (GSEA) of ranked gene expression changes confirmed these findings. Synaptic and signaling-related gene sets showed positive normalized enrichment scores (NES), indicating enrichment among up-regulated genes (**Figure 2E**). In contrast, immune-related gene sets displayed negative NES values, indicating enrichment among down-regulated genes (**Figure 2F**). These results suggest that GC exposure in the ACC modulates transcriptional programs toward neuronal and synaptic functions while suppressing immune signaling.

**Figure 2.**
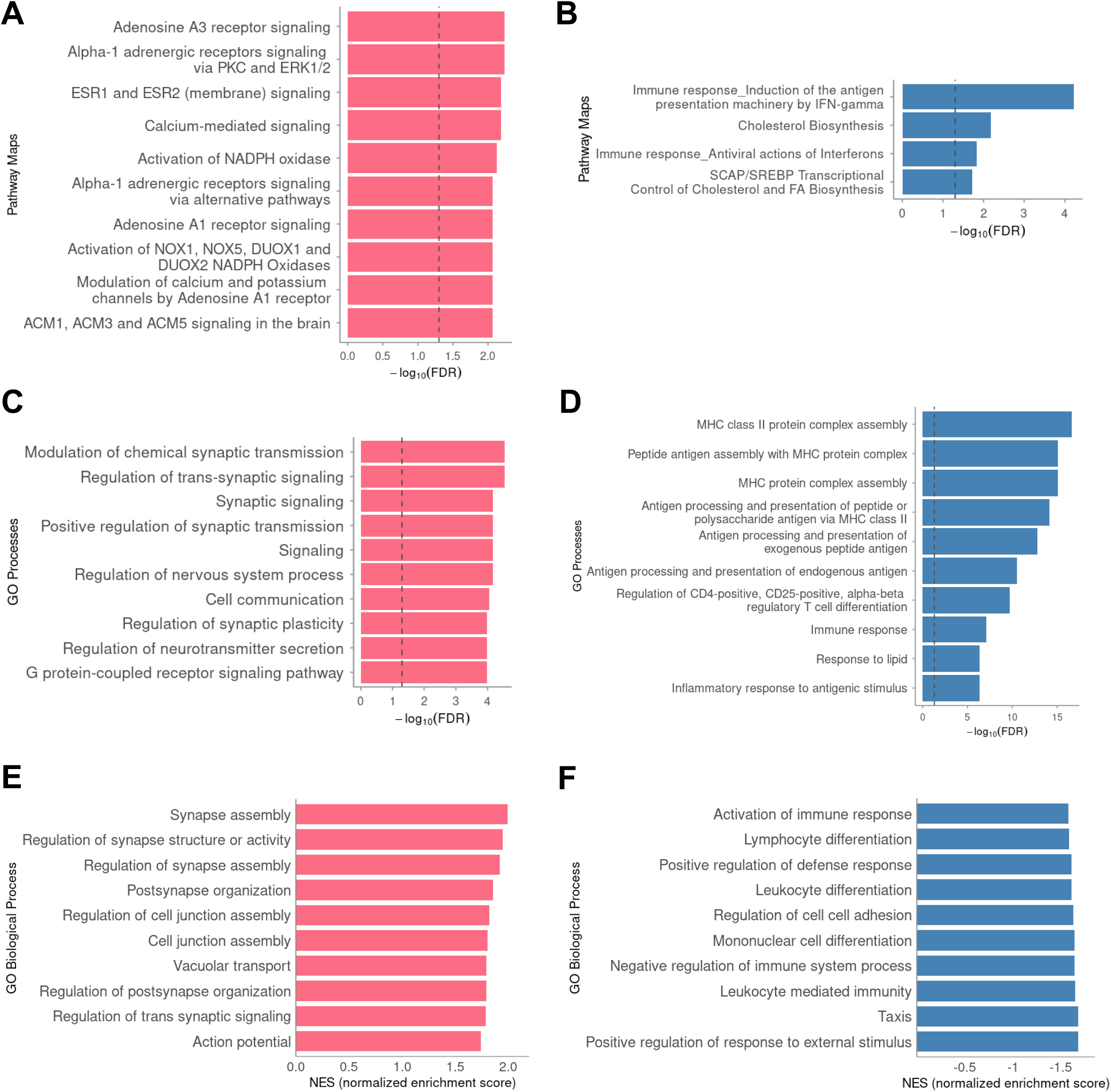
Functional enrichment analysis of differentially expressed genes (DEGs). MetaCore over-representation analysis of enriched Pathway Maps (**A–B**) and Gene Ontology (GO) Biological Processes (**C-D**) among DEGs (FDR< 0.05). Panels (**A**) and (**C**) show enrichments for upregulated genes, whereas panels (**B**) and (**D**) correspond to downregulated genes. Bars represent the –log₁₀(FDR) for the top significantly enriched categories identified using MetaCore (Clarivate Analytics). Gene set enrichment analysis (GSEA) of GO Biological Processes is shown for upregulated (**E**) and downregulated (**F**) genes, with bars indicating normalized enrichment scores (NES). Differential expression was assessed using RNA-seq data from anterior cingulate cortex (ACC) tissue, with statistical significance defined at FDR < 0.05.

### Network-level transcriptional responses to glucocorticoids

We applied a weighted gene co-expression network analysis (WGCNA) ^42^ to determine GC-induced alterations across the ACC transcriptional architecture. WGCNA computes pairwise correlations between gene expression profiles to reveal clusters of highly co-expressed gene modules based on shared patterns of expression reflecting gene networks ^50^. Module eigengenes were correlated with the experimental condition to identify the most GC-responsive networks ^51^. WGCNA identified 18 co-expression modules in the ACC data set (**Figures 3A and 3B**). The Red (r=0.61, p=0.04), Blue (r=0.66, p=0.02), and Green (r=0.77, p=0.003) modules showed significant positive correlations with GC exposure, indicating networks with coordinated transcriptional responses to betamethasone. Importantly, both Blue and Green but not the Red module were also significantly enriched for the GC-induced DEGs (Fisher’s exact test, Blue FDR= 1.32x10^-9^ and Green FDR= 0.003, **Figure 3B**), supporting their biological relevance as GC-regulated networks. Subsequent analyses focused on the Blue and Green modules.

**Figure 3.**
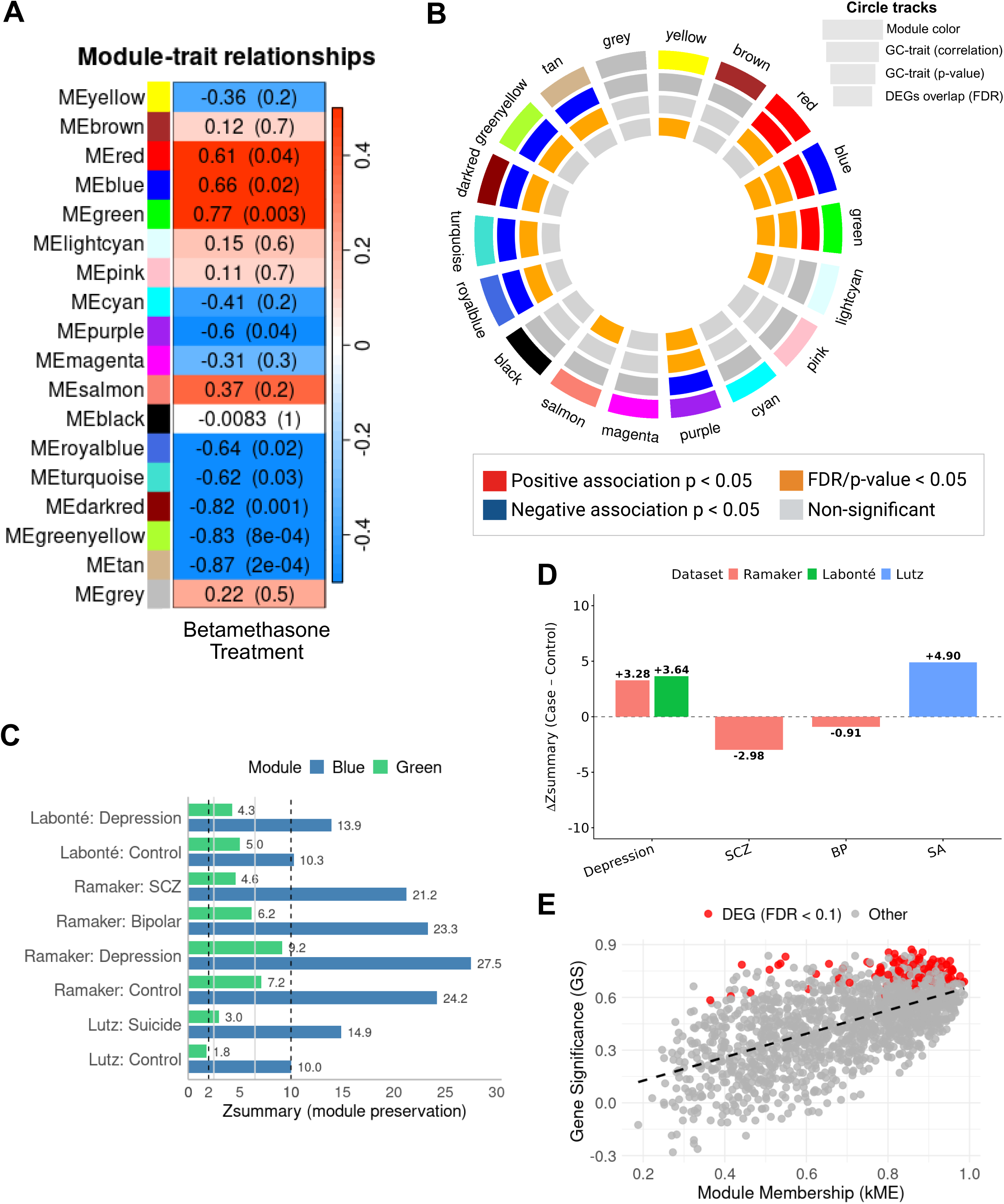
WGCNA module–trait associations and module preservation. **(A)** Heatmap of module–trait relationships showing correlations between WGCNA module eigengenes and betamethasone treatment in the macaque anterior cingulate cortex (ACC). Correlation coefficients are shown, with corresponding p-values in parentheses. **(B)** Circular plot summarizing module-level properties, including module color assignment, direction and significance of glucocorticoid (GC)–trait correlations, and overlap with differentially expressed genes (DEGs). **(C)** Preservation of selected modules across independent human postmortem ACC datasets (Ramaker et al., 2017; Labonté et al., 2017; and Lutz et al., 2017), quantified using Zsummary statistics (also see **Figures S1-4**). **(D)** Differential module preservation expressed as ΔZsummary (case-control) for psychiatric conditions across datasets. **(E)** Relationship between module membership (kME) and gene significance (GS) within the Blue module, with differentially expressed genes (FDR < 0.1) highlighted in red.

### Preservation of GC-responsive co-expression networks in human brain

We performed module preservation analyses using the WGCNA framework ^52^ to evaluate whether the GC-responsive co-expression networks in macaque ACC reflect conserved transcriptional networks in human brain. This approach was essential for the translation of the molecular signal from non-human primate samples into analyses with human data sets. Module preservation assesses whether the topological properties of a module (intramodular connectivity, density, and overall structural organization) are reproducible in independent datasets ^52^.

Preservation statistics (Zsummary and MedianRank) were computed using permutation-based tests across three independent human post-mortem RNAseq datasets, including clinical and matched control samples. Across all datasets, the Blue GC-responsive network showed a remarkably consistent and significant preservation in both clinical and control samples (**Figure 3C**). Analysis using the ACC dataset from Ramaker et al.^53^ showed that the Blue module was strongly preserved in depression (Zsummary = 27.5), bipolar disorder (Zsummary = 23.3), and schizophrenia (Zsummary = 21.2), as well as healthy controls (Zsummary = 24.2). Similar results were observed in the ACC dataset from Lutz et al. ^54^, with high preservation in suicide (Zsummary = 14.9) and borderline-high preservation in controls (Zsummary = 10.0). The cingulate gyrus dataset from Labonté et al. ^55^ confirmed these findings, with the module preserved in both MDD (Zsummary = 13.9) and controls (Zsummary = 10.3) (full Zsummary and MedianRank results are depicted in **Figures S1-S4)**. In contrast, the Green showed only moderate preservation across most datasets, except in Lutz et al ^54^ control group (**Figure 3C**), where preservation was not detected, suggesting that this module is notably less consistently conserved in human ACC than is the Blue module. Finally, to compare preservation strength directly between diagnostic groups, we computed ΔZsummary (disease - control) for the Blue module (**Figure 3D**). Depression and suicide conditions showed positive ΔZsummary values, indicating stronger preservation in affected individuals relative to controls, whereas schizophrenia and bipolar disorder displayed negative ΔZsummary values. Together, these results demonstrated that the Blue module represents a highly conserved, GC-regulated transcriptional network in the human cingulate cortex RNAseq data sets and notably those from MDD and suicide samples.

### Refinement of GC-responsive modules for translational analyses

The Blue ACC gene network was then refined to enhance biological specificity for translational analysis in humans by intersection with the set of GC-responsive DEGs (FDR<0.1). This filtering step isolated genes that were both co-expressed within the network and directly regulated by GC exposure, while reducing noise introduced by the large size of the original module (1526 genes). This refinement yielded a core set of 107 genes constituting the refined Blue module (**Figure 3E**) and ensured that analyses in human data sets focused on the components most likely to mediate the GC-related signal. To then enable translational analyses in human cohorts, orthologous human genes corresponding to the refined Blue module were identified using the Ensembl biomaRt (GRCh37.p13)^56^. This mapping reduced the refined Blue module from 107 macaque genes to 96 human orthologs retained for downstream polygenic score analyses (gene list provided in **Table S2**).

### Translation of preserved GC-sensitive networks in humans

The strong preservation of Blue gene network suggests a GC-transcriptional architecture is conserved in humans and may reflect inter-individual differences in capacity for GC-induced effects on gene expression in the ACC and thus increased sensitivity to adversity. To translate these findings to humans, we employed a bioinformatic approach to interrogate human datasets enabling tests of specific hypotheses concerning the molecular mechanisms that link adversity to specific mental health outcomes through gene x environment analyses in the UK Biobank (baseline characteristics of the study sample are summarized in **Table 1**). We focused on the moderating role of GC signaling in the ACC, a region closely associated with depression. Genes from the revised ACC Blue gene network were used as the basis for the calculation of an expression-based polygenic score (ePGS) in humans (see Methods) derived from transcriptomic data. An ePGS score is a cumulative index of functional single-nucleotide polymorphisms (SNPs) that are associated in cis with the expression of the genes in a target gene network (i.e., an expression quantitative trait locus; eQTL; see **Figure 4A**). A functional SNP was defined as a common variant statistically associated with the level of expression for any gene in the ACC Blue gene network using the Genotype-Tissue Expression (GTEx; https://gtexportal.org/home/) database that includes both genotyping and brain region-specific RNAseq. An ePGS thus reflects variation in the genetically determined capacity for the transcriptional activity of the network genes: A higher ePGS reflects a greater capacity for GC-induced expression in the ACC of genes comprising the Blue module network. According to our hypothesis, this ePGS should then predict individual differences in mental health outcomes as a function of adversity as reflected in gene x environment interaction effects.

**Figure 4.**
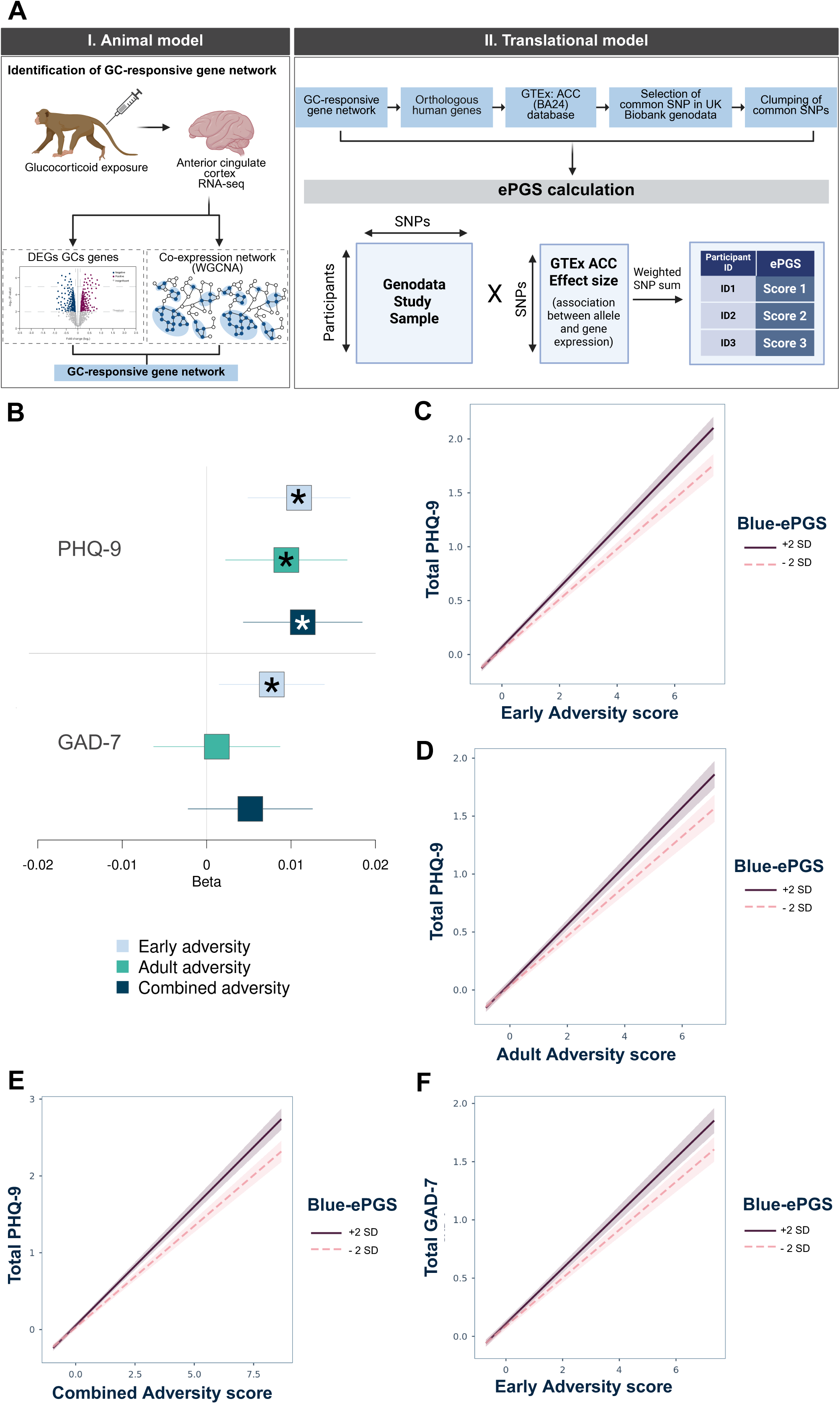
Derivation of a glucocorticoid-responsive polygenic score and its interaction effect with adversity on depressive and anxiety symptoms. **(A)** Translational framework for expression-based polygenic score (ePGS) construction. Glucocorticoid (GC)-responsive gene networks were first identified in the anterior cingulate cortex (ACC) of non-human primates exposed to GC using RNA-sequencing (RNA-seq). Differential expression and weighted gene co-expression network analyses (WGCNA) were performed to identify gene modules associated with GC treatment. From the GC-associated module showing a positive relationship with treatment, genes were further filtered to retain only those differentially expressed at FDR < 0.1, thereby restricting the network to genes both co-expressed and transcriptionally responsive to GC exposure. Human orthologs of the filtered gene set were then overlapped with ACC cis-eQTLs derived from the Genotype-Tissue Expression (GTEx) project and UK Biobank. Common SNPs were LD-clumped to obtain a non-redundant variant set and weighted by GTEx-estimated expression effect sizes to generate the ePGS. **(B)** Standardized gene-environment interaction effects (β ± SE) between the Blue GC-responsive ePGS/ACC and early, adult, and combined adversity for depressive (PHQ-9) and anxiety (GAD-7) symptoms in UK Biobank females. Asterisks indicate significant interaction effects after FDR correction (FDR<0.05). (**C-E**) Simple slopes for significant interaction effects between the Blue ePGS/ACC and (**C**) early life, (**D**) adult, and (**E**) combined adversity in relation to depressive symptoms (PHQ-9). **(F)** Simple slopes for significant interaction between refined Blue ePGS/ACC and early life adversity in relation to anxiety symptoms (GAD-7). Higher adversity exposure is associated with increases in depressive/anxiety symptoms at higher ePGS levels. Shaded areas represent 95% confidence intervals. Interaction effects were estimated using linear regression models adjusted for relevant covariates.

**Table 1.** Baseline characteristics of the study sample stratified by Blue module ePGS/ACC. N = 81,854 (participants with either PHQ-9 or GAD-7 and either early or adult adversity data available).

| Variable | Lower ePGS | Higher ePGS | p-value |
| --- | --- | --- | --- |
| **Demographics** |  |  |  |
| Age, years | 55.13 ± 7.58 | 55.26 ± 7.62 | 0.013 |
| Townsend deprivation index | -1.63 ± 2.88 | -1.68 ± 2.83 | 0.008 |
| **Genetic ancestry, N (%)** |  |  | < 1 × 10<sup>-8</sup> |
| European ancestry | 33,437 (82.8) | 34,960 (84.3) |  |
| Non-European ancestry | 6,945 (17.2) | 6,512 (15.7) |  |
| **Lifestyle factors** |  |  |  |
| Smoking status, N (%) |  |  | 0.20 |
| Never | 24,800 (62.4) | 25,247 (62.6) |  |
| Previous | 12,897 (32.4) | 13,490 (33.5) |  |
| Current | 2,593 (6.5) | 2,648 (6.6) |  |
| **Clinical measures** |  |  |  |
| Depression diagnosis, N (%) |  |  | 0.17 |
| No | 22,439 (74.4) | 22,938 (74.9) |  |
| Yes | 7,721 (25.6) | 7,693 (25.1) |  |
| GAD-7 total score | 2.44 ± 3.57 | 2.45 ± 3.58 | 0.72 |
| PHQ-9 total score | 3.03 ± 3.82 | 3.06 ± 3.88 | 0.33 |
| **Adversity measures** |  |  |  |
| Early-life adversity | 1.88 ± 2.70 | 1.83 ± 2.64 | 0.009 |
| Adult adversity score | 2.44 ± 2.81 | 2.40 ± 2.82 | 0.17 |
| Combined adversity score | 4.32 ± 4.48 | 4.24 ± 4.47 | 0.036 |

Values are presented as mean ± SD for continuous variables and N (%) for categorical variables.
Continuous variables were compared using two-sided t-tests and categorical variables using Pearson's chi-squared tests. P-values are unadjusted and are provided for descriptive purposes only.

Importantly, the Blue network ePGS scores showed no significant main effect for association with self-reported symptom measures of depression (Patient Health Questionnaire-9; PHQ-9) and anxiety (Generalized Anxiety Disorders-7; GAD-7) in the UK Biobank (**Table S3)**. We hypothesized significant gene x environment interactions using the Blue module ePGS/ACC and measures of exposure to adversity specific to early childhood, adulthood or their cumulative index (see Methods). Importantly, we showed that all three adversity scores presented robust main effects on the PHQ-9 and GAD-7 outcomes (p<0.0001, **Table S3**). This finding is consistent with the established role of adversity as an environmental risk factor and enables a valid test of the GC moderation hypothesis. We then tested whether the ePGS moderated the effect of adversity exposure on self-reported symptoms of depression and anxiety.

Analysis of data from adult females in the UK Biobank revealed significant interaction effects between the Blue module ePGS/ACC and early-life adversity (FDR=0.002, β=0.011, N=80877), adult adversity (FDR=0.020, β=0.009, N=57888), and the combined adversity score (FDR=0.005, β=0.011, N=57312) on PHQ-9 symptoms (**Figure 4B, Table S4)**. No significant interactions were detected in males (**Table S5**), revealing sex-specificity. Although interactions were observed for multiple adversity measures, the combined adversity score did not exceed the magnitude of the individual effects. Instead, the interaction with early-life adversity showed the strongest and most consistent effect, suggesting that the Blue module ePGS/ACC preferentially captures sensitivity to early stress exposure.

Simple slope analyses were performed to further characterize the gene x environment interactions observed in females. Note, ePGS score is comprised of SNPs known to increase the expression of the network genes in the human ACC and is thus assumed to reflect a greater capacity for the expression of genes in the GC-sensitive Blue network. The results revealed that, as predicted, individuals with higher Blue module ePGS/ACC values showed more pronounced increases in depressive symptoms as a function of exposure to adversity. This pattern was observed for early-life adversity (higher-ePGS: β = 0.27, p < 0.0001; lower-ePGS: β = 0.24, p < 0.0001; **Figure 4C**), adult adversity (higher-ePGS: β = 0.4, p < 0.0001; lower-ePGS: β = 0.22, p < 0.0001; **Figure 4D**) and the combined adversity score (higher-ePGS: β = 0.30, p < 0.0001; lower-ePGS: β = 0.28, p < 0.0001; **Figure 4E**).

Given the relation between depression and anxiety, we examined anxiety symptoms measured by the GAD-7 in adult females. A significant interaction effect was observed between the Blue ePGS/ACC and early-life adversity (FDR= 0.02, β= 0.008, N= 81120, **Figure 4F, Table S4**) but not measures of adult or combined adversity. Simple slope analyses revealed that individuals with higher Blue-ePGS/ACC values showed greater increases in anxiety symptoms with increasing early-life adversity exposure (higher-ePGS: β = 0.23, p < 0.0001; lower-ePGS: β = 0.21, p < 0.0001; **Figure 4F**).

We used the ePGS pipeline to assess the biological specificity of the observed gene x environment interactions in females of the Blue module ePGS/ACC by generating ePGS scores for the Green and Tan modules, the latter of which was very significantly and negatively associated with GC exposure (r=-0.87, see **Figure 3A**). The modules were filtered for ePGS calculation as described above for the Blue module. Neither the ePGS scores based on the refined Green or Tan modules showed evidence of interaction effect with early, adult, or combined adversity on either PHQ-9 or GAD-7 outcomes in adult females (all interaction effects p ≥ 0.10, **Table S6**).

An important challenge in the study of MDD is that of identifying the neurobiological basis for specific depressive symptoms. Item-level analyses of the individual PHQ-9 symptoms in females (**Figure 5, Table S7**) revealed that interactions between the refined Blue-ePGS/ACC and adversity were most prominent for core cognitive–affective symptoms, consistent with the known role of the ACC in brain health. Significant interactions involving the Blue-ePGS/ACC were apparent across all three adversity measures for suicidal ideation and concentration difficulties, while early-life and combined adversity were significantly associated with feelings of inadequacy and depressed mood. In contrast, the interaction of the Blue-ePGS/ACC with adult and combined adversity was significant only for anhedonia. Suicidal ideation emerged as the most consistent item across adversity exposures, with early-life adversity showing the strongest statistical evidence (p= 6.7 x 10^-5^, β =0.012), and broadly comparable effect sizes observed for combined adversity (p= 2.0 x 10^-4^, β =0.013).

**Figure 5.**
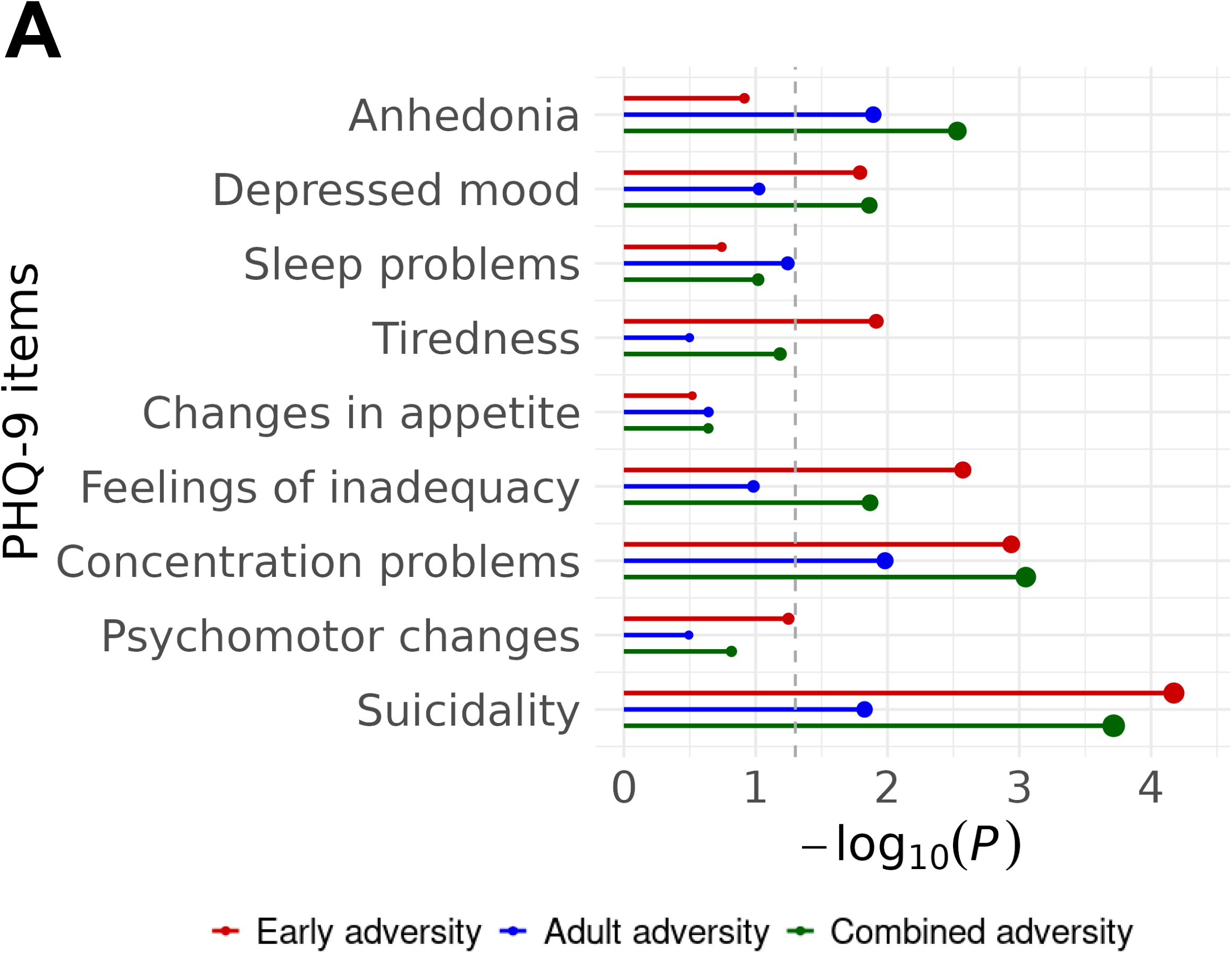
Symptom-level gene-environment interaction effects. Gene–environment interaction effects between the refined Blue GC-responsive ePGS and adversity scores at the symptom-item level for depressive symptoms (PHQ-9) in UK Biobank females. Each point represents the −log₁₀(P) value of the interaction between the ePGS and early-life, adult, or combined adversity for a given symptom item. The dashed vertical line indicates nominal statistical significance (P = 0.05).

### Developmental and clinical variation patterns in the Blue module co-expression

The moderation effect of the Blue module ePGS/ACC was stronger and more consistent with measures of early life adversity. Accordingly, we investigated whether patterns of gene co-expression within this network vary across human developmental stages. Pairwise expression correlations among Blue module ePGS/ACC genes (96 genes) were performed using BrainSpan human brain expression data that includes brain region-specific transcriptomic data in childhood and adulthood.

Across all pairwise gene–gene expression correlations within the module, transcriptomic data from children (**Figure 6A**) showed markedly stronger co-expression than that from adults (**Figure 6B**), as reflected by higher mean and median correlation coefficients and a substantially greater proportion of strongly correlated gene pairs (|r| > 0.5; OR = 3.99, 95% CI = 3.65–4.37; Fisher’s exact p = 2.46 × 10⁻²¹⁷, **Figure S5A**). Group differences were highly significant based on distributional comparisons (Wilcoxon test, W = 5.65 × 10⁶, p = 1.19 × 10⁻²⁷⁴) and were further confirmed using permutation-based testing (Δ = 0.21, p_perm_ < 1 × 10⁻⁴). In sum, the expression of the genes comprising the Blue module ePGS/ACC showed stronger inter-correlations in childhood than in adulthood, suggesting a greater coherence of the Blue module gene network during development.

**Figure 6.**
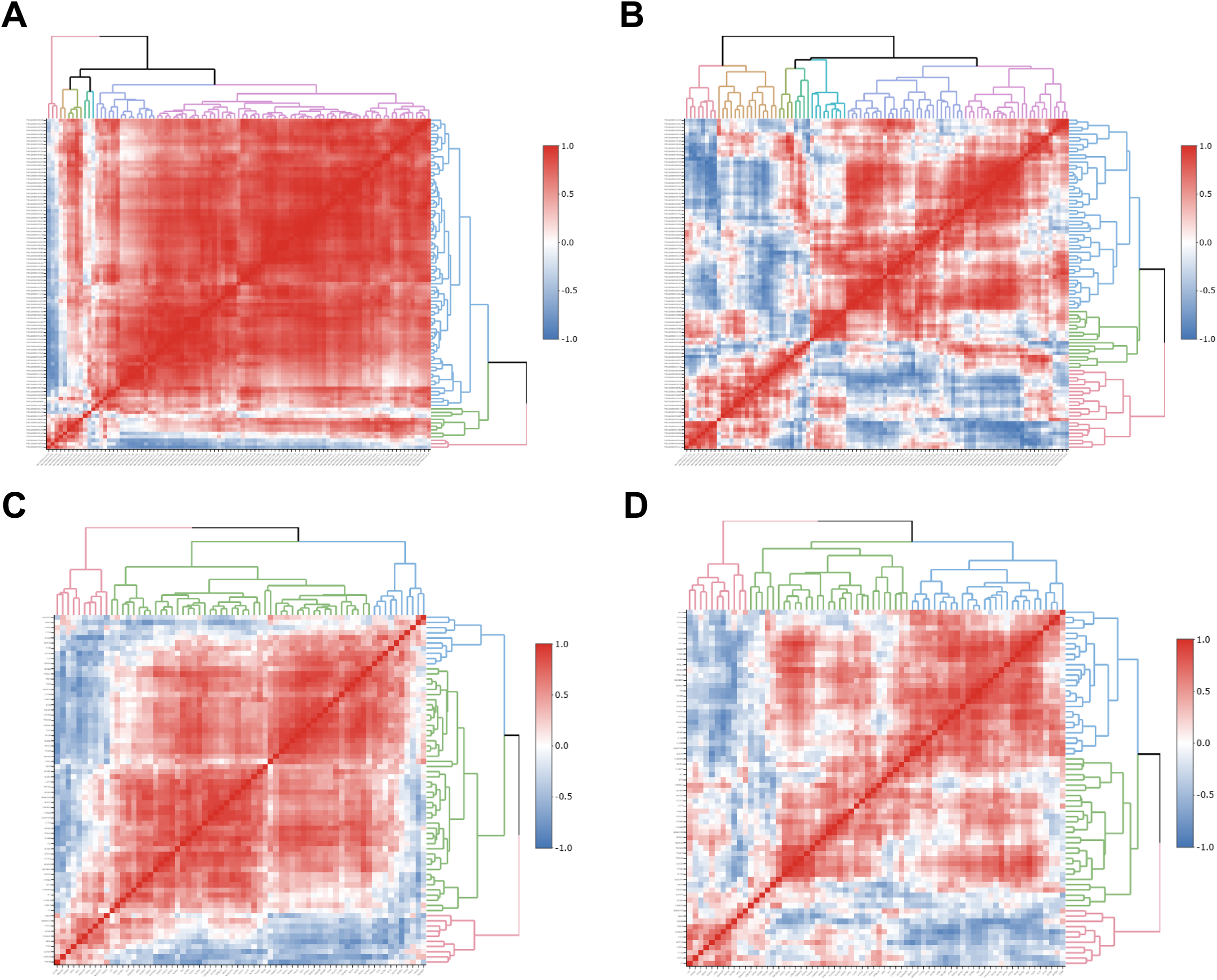
Human co-expression patterns of the Blue module ePGS/ACC network. Heatmaps illustrating gene-gene expression patterns of the genes comprising the refined Blue-ePGS network across independent cingulate cortex transcriptomic datasets. Gene-gene correlation matrices are shown for **(A)** childhood samples from BrainSpan (ages 1-13, N=7), **(B)** adulthood samples from BrainSpan (ages 19-37, N=6), **(D)** control subjects from Lutz et al. 2017 (N=23), and **(E)** suicide cases with history of severe childhood abuse from Lutz et al. 2017 (N=27). Rows and columns correspond to genes within the refined Blue network, ordered by hierarchical clustering. Color scale indicates the strength and direction of correlations (red, positive; blue, negative). Also see **Figure S5**.

We then used the same post-mortem dataset previously employed for module preservation analyses ^54^ to examine co-expression of the Blue module genes in a clinical context. Compared with controls, suicide cases with history of child abuse exhibited significantly higher pairwise gene–gene correlations within the genes identified in the refined Blue module ePGS (Wilcoxon test, W = 3.05 × 10⁶, p = 1.84 × 10⁻²⁵), alongside an increased proportion of strongly co-expressed gene pairs (|r| > 0.5; OR = 1.70, 95% CI = 1.50–1.92; Fisher’s exact p = 9.27 × 10⁻¹⁸, **Figure S5B**). This pattern was further supported by permutation testing (Δ = 0.074, p_perm_ < 1 × 10⁻⁴; **Figure 6C** and **6D** for suicide and controls, respectively). These findings taken together indicate that the genes that comprise the refined Blue module ePGS/ACC exhibit developmentally and clinically sensitive co-expression patterns in humans, consistent with the interaction effects observed between refined Blue-ePGS and early-life adversity on symptoms of anxiety and depression.

### Functional characterization of GC-responsive Blue ePGS gene network

We examined topological organization and functional annotations to better characterize the biological relevance of the 96 human ortholog genes in the refined GC-responsive Blue ePGS network. Network centrality analyses identified highly connected genes with potential regulatory importance. Centrality analyses showed *GFRA1*, *FOXN3*, and *DCLK1* as the top hub genes within the refined Blue module sub-network (**Figure 7A** and **7B**). These genes converge on biological processes related to neurodevelopment, neuronal maintenance and transcriptional regulation. *GFRA1* encodes a co-receptor for glial cell line–derived neurotrophic factor (GDNF), a key mediator of neuronal survival, differentiation, and plasticity. Studies of humans have associated *GFRA1* with depression-related phenotypes, including suicidal ideation and behavior ^57–59^. *DCLK1* (doublecortin-like kinase 1) is involved in neuronal migration, cytoskeletal organization, and apoptosis, processes critical for brain development and structural plasticity.

**Figure 7.**
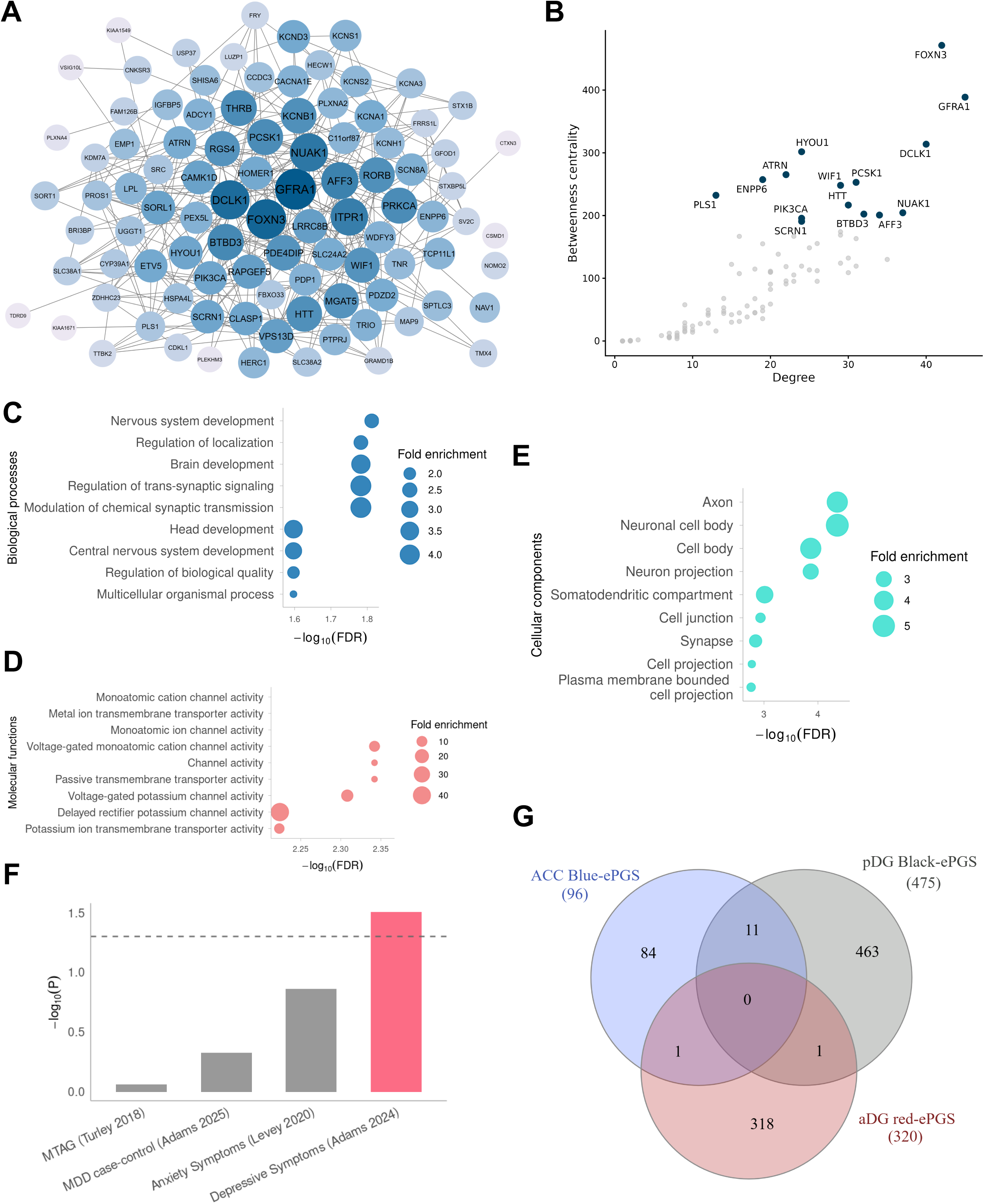
Network topology, functional enrichment, and genetic relevance of the Blue module ePGS/ACC gene network. **(A)** Network representation of the Blue module ePGS gene set visualized in Cytoscape. Node color intensity reflects genes with higher degree centrality. **(B)** Scatter plot showing the relationship between node degree and betweenness centrality for genes in the Blue ePGS network, derived from Cytoscape/Centiscape network analysis. Highly central hub genes are labeled. **(C–E)** Gene Ontology enrichment analysis of the Blue ePGS network, showing enriched Biological Processes **(C),** Molecular Functions **(D)**, and Cellular Components **(E)**. Point size reflects fold enrichment, and the x-axis indicates −log₁₀(FDR). **(F)** Enrichment of the Blue ePGS gene set in genome-wide association study (GWAS) summary statistics assessed using MAGMA. Bars represent −log₁₀(P) values for enrichment across neuropsychiatric traits, with the dashed line indicating nominal statistical significance. (**G**) Venn diagram showing minimal gene overlap between the Blue module ePGS/ACC network and GC-responsive gene networks previously identified in the hippocampal dentate gyrus.

*FOXN3*, a member of the forkhead box family of transcription factors, binds to DNA to control gene expression. Polygenic analyses have implicated *FOXN3* among neurodevelopmental genes associated with suicide attempt ^60^. Together, these hub genes highlight a network centered on developmental and plasticity-related mechanisms, consistent with the sensitivity of the Blue module gene network to early-life adversity and stress-related depressive phenotypes, especially suicidal ideation. Consistent with this interpretation, a functional enrichment analysis demonstrated that genes within the refined Blue gene network were primarily involved in nervous system development and synaptic signaling (**Figure 7C**). Molecular function terms were strongly enriched for ion channel and transporter activity (**Figure 7D**), while cellular component annotations highlighted localization to the neuronal cell body, axon, and synapse (**Figure 7E**).

These functional profiles align with the identity of the top hub genes and collectively indicate a synaptic and neurodevelopmental signature. Finally, GWAS enrichment analysis using MAGMA revealed nominal associations between the refined Blue network genes and GWAS of depressive symptoms (**Figure 7F**). In sum, these analyses are consistent with the relevance of the GC-responsive Blue gene network of the ACC in moderating the impact of early-life adversity on the risk for MDD, especially suicidal ideation.

To place these functional and topological features in a broader anatomical context, we compared the refined ACC Blue ePGS gene network with GC-sensitive gene networks previously identified in the posterior and anterior hippocampal dentate gyrus in the same female macaques ^47^. Overlap analysis revealed minimal gene sharing across regions, with 11 genes shared between the ACC Blue and posterior dentate gyrus networks, one gene shared with the anterior dentate gyrus network, and no genes common to all three networks (**Figure 7G**), indicating notable tissue-specificity of the ACC-specific GC-responsive transcriptional program.

## DISCUSSION

The spatial and temporal heterogeneity of gene expression provides a major challenge for understanding molecular mechanisms underlying the risk for psychopathology ^39^. Transcriptomic data sets, unlike those of genotype, offer spatial resolution and informatic analyses enable the identification of gene networks. Transcriptomic analyses also offer the opportunity to document alterations in gene expression in response to conditions of established relevance for brain health. Accordingly, we characterized the transcriptional sensitivity to chronically elevated GC exposure, focusing on the ACC using a non-human primate model (**Figure 1A**). The focus was based on the known relation between increased GC exposure and mental health, as well as the importance of the ACC, especially for MDD and suicidal ideation. The validity of the model was reflected in the altered expression of genes such as *FKBP5*, a well-established GC target and closely associated with psychopathology ^61–63^ as well as informatic analyses revealing that GC-induced effects on gene expression were associated with well-established GC-regulated processes such as immune activation, noradrenergic signaling and insulin sensitivity.

WGCNA revealed three modules significantly associated with chronic GC exposure. Of these, the Blue module demonstrated a rather remarkable level of preservation when compared with a range of human post-mortem RNAseq data sets (**Figure 3C**). Each of these comparison data sets bear clinical relevance for MDD, including post-mortem human ACC from MDD and suicide samples. We thus focused on the Blue module for the calculation of a transcriptomic-based ePGS, limiting inclusion to those genes showing both high centrality and significantly GC-induced altered ACC expression. This gene network approach is consistent with the established polygenic definition of vulnerability for MDD ^64,65^. The Blue module ePGS/ACC was then used to test the hypothesis that increased capacity for the expression of GR-regulated genes in the ACC would render individuals more sensitive to the effects of adversity. We selected the PHQ-9 as the critical outcome measure for the analyses using the UK Biobank. The advantages of this approach derived from the continuous nature of the measure, as opposed to categorical ICD diagnosis, as well as the ability to subsequently perform a secondary analysis focusing on individual depressive symptoms (**Figure 5**). The Blue module ePGS significantly moderated the association between adversity and PHQ-9 scores, a finding consistent with our hypothesized role for GC sensitivity.

A striking finding in the Blue ePGS x adversity analyses with the total PHQ-9 score was the notably stronger moderation of the association with early-life adversity compared to recent adversity in adulthood (**Figure 4B, Table S4**). This pattern, with one notable exception, was also apparent in the secondary analysis performed using individual PHQ-9 items. The scores for depressed mood, tiredness, concentration problems, feelings of inadequacy, and suicidality, the significance of the moderating effect of the Blue ePGS was greatest for childhood adversity (**Figure 5, Table S7**). The analysis using the GAD-7 revealed a significant moderating effect of the Blue ePGS only for childhood adversity. These findings suggest that an ACC-specific, GC-responsive gene network influences susceptibility to the effects of early-life adversity on the risk for depression over the lifespan.

The more prominent moderating effect of the Blue ePGS/ACC for early-life adversity is consistent with the results of analysis of the underlying gene network (**Figures 6A-B**) using the BrainSpan human gene expression database. The heatmap plot for gene–gene co-expression showed greater widespread co-expression in childhood (ages 1 to 13 years) than in adulthood. The number of co-expressed genes (i.e., with correlations of r> 0.5) in the Blue module was more than twice that in childhood compared with samples from adults. This finding suggests the GC-regulated Blue module gene network is more coherent in early life, consistent with the finding of a greater moderating effect of childhood adversity. This finding aligns with evidence favoring the role for GC signaling during early development in shaping the risk for later psychopathology ^16,66,67^. Genetic variants that significantly moderate the response to GC treatment and are enriched for those associated with psychopathology, as well as for increased expression across neurodevelopment ^18^. This developmental enrichment, together with the established role of GCs as mediators of the effects of stressors, suggested that GR-regulated gene networks might mediate the effects of early-life adversity on later psychopathology ^41^. In this context, our results suggest that a subset of genetic variants associated with the risk for MDD acts through a stress-induced increase in activity of brain region-specific, GR-regulated gene networks. These findings are consistent with those of broader genomic analyses showing that genes associated with psychiatric disorders are enriched for expression in perinatal development and associated with neurodevelopmental processes ^39,64,68^.

The present results with the Blue ePGS/ACC contrast with those previously reported for analyses of transcriptomic analyses of the anterior and posterior hippocampal dentate gyrus ^47^ using tissues from the same betamethasone- and saline-treated animals. The GC-regulated transcriptome shows remarkable variation across brain regions ^16,25^. RNAseq analyses using our non-human primate model likewise reveal across the anterior and posterior hippocampal dentate gyrus and ACC. The modules identified using WGCNA as those most significantly associated with chronic GC treatment showed literally no overlapping genes between them (**Figure 7G**).

Likewise, the moderation of the effect of adversity by the ePGS scores derived from these modules associated with very different mental health outcomes. In the hippocampus, the ePGS score derived from the module most strongly associated with GC treatment significantly moderated the impact of adversity on psychotic states ^47^. There was no effect on depression or anxiety. In contrast, the ePGS derived from the GC-responsive Blue module in ACC significantly moderated the effect of adversity on depression and anxiety, but not psychotic states (data not shown). Extensive analyses of large human clinical data sets revealed effects of prolonged GC treatment on the risk for depression, anxiety and suicide as well as psychosis ^7–10^. These findings suggest that GC effects on specific mental health outcomes are mediated through actions at very different brain regions. Moreover, since ePGS scores are based in part on genetic variants associated with region-specific regulation of GC-responsive genes, the mental health outcomes associated with prolonged GC exposure may vary as a function of genotype. While speculative, this suggestion is consistent with the results of Arloth et al. ^18^, who showed that genetic variants that significantly moderated the transcriptional response to a GR agonist were enriched among risk alleles for psychiatric disorders.

Analyses using the UK Biobank data of self-reported symptoms of depression showed that the moderating influence of the Blue ePGS/ACC on the effects of adversity were unique to females. This finding is likely related to the fact that the transcriptomic data from which the ePGS was devised were derived from female macaques (males were not available for these experiments at the facility). Moreover, recent sex-stratified genome-wide association studies of MDD reveal sex differences in the relevant genetic architecture ^69,70^. Likewise, transcriptomic analyses of MDD using human post-mortem brain reveal a remarkable degree of sex-specificity, particularly in the PFC ^55,71–76^.

The transcriptomic approach to the calculation of polygenic scores provides an advantage of tissue-specificity lacking in polygenic risk scores derived from genotype. Moreover, the experimental design of the current study identified GC-responsive genes and thus created a polygenic score specific to a clinically-relevant physiological condition. Nevertheless, our analyses bear limitations. Transcriptomic data included only female macaques due to limitations related to our colony and thus did not allow for analysis of sex-specific polygenic scores.

Another limitation is that our study was based on bulk RNA-sequencing, which merges contributions from composite cell types. Future studies using single-cell RNA-sequencing will provide cell-type-specific transcriptional profiles to further refine the development of transcriptomic-based polygenic scores. The exclusive use of European cohorts in this study is a limitation because the results may not necessarily generalize to other populations. These limitations notwithstanding, our study provided novel insights into the tissue-specific mechanisms of chronic GC exposure in defining susceptibility to adversity for later psychopathology.

## ACKNOWLEDGEMENTS

We acknowledge the UK Biobank participants and staff for their invaluable contribution to this research. This research has been conducted using the UK Biobank Resource under application number 41975. This study was funded by a grant from the Hope for Depression Research Foundation (MJM). Funding for the non-human primate colony was provided by the National Medical Research Council of Singapore.

## DECLARATION OF INTERESTS

The authors declare no competing interests.

## METHODS

### EXPERIMENTAL MODEL AND STUDY PARTICIPANT DETAILS

#### Animal model

Adult female macaques (*Macaca fascicularis,* 7-20 years) were housed singly or in pairs under controlled environmental conditions at the SICS large animal facility (SMEC, Singapore), with ad libitum access to water and a balanced commercial diet enriched with fresh fruits. All experimental procedures were approved by the Singapore Health Institutional Animal Care and Use Committee (IACUC; protocols #2014/SHS/1016 and #2015/SHS/1078). Experiments adhered to the Guidelines on the Care and Use of Animals for Scientific Purposes (National Advisory Committee for Laboratory Animal Research - NACLAR, Singapore), aligned with Australian Code of Practice for the Care and Use of Animals for Scientific Purposes (NHMRC, Australia), Guide for the Care and Use of Laboratory Animals (National Academy of Sciences, USA), and The Good Practice Guide for the Use of Animals in Research (New Zealand). Trained veterinarians performed all interventions.

Animals were randomly assigned to two groups of similar age and received daily intramuscular injections of saline (n = 6) or betamethasone (Diprosan, 0.3 mg/kg/day; n = 6) for seven consecutive days. Only females were included, as they were the available subjects from the facility’s breeding program at the time of the study. The physiological efficacy of chronic betamethasone administration in this cohort has been previously established, with circulating cortisol levels reduced by more than 90% relative to controls ^47^. Thus, circulating cortisol levels following betamethasone treatment were reduced by more than 90% of control values (Mean ± SD: 25.9 ± 6.5 vs. 1.8 ± 1.7 mg/dL; p<.0001), reflecting the expected negative-feedback effect of the GR agonist and the physiological efficacy of the treatment.

#### Human participants

The UK Biobank is a large population-based cohort from the United Kingdom^77^. Participants were recruited through assessment centers between 2006 and 2010 (ages 37-73), resulting in a total of 502,543 individuals. The study received ethical approval from the North West Multicentre Research Ethics Committee (REC reference 11/NW/0382), the National Information Governance Board for Health and Social Care, and the Community Health Index Advisory Group. Analyses were conducted under UK Biobank application number 41975. All participants provided informed consent and completed baseline assessments, which included questionnaires, physical measurements, and the collection of biological samples.

Phenotypic information is available for 502,543 individuals, of whom 487,707 have genome-wide genotyping data. Analyses were restricted to unrelated individuals and excluded participants who withdrew consent, lacked genotyping data, showed inconsistencies between genetic and self-reported sex, or were heterozygosity outliers. Participants eligible for inclusion were those with available genotyping data, the phenotypic outcome under investigation, and valid environmental information to compute the environmental scores. After applying these criteria, the final analytic sample comprised 146,162 participants (81,854 females and 64,308 males) as depicted in **Figure S6**. Descriptive characteristics of the study sample are presented in **Table 1**.

## METHOD DETAILS

### Tissue collection and RNA-sequencing

On day eight (∼10 AM), animals were euthanized by intramuscular ketamine injection (7–10 mg/kg) followed by intravenous pentobarbital (80–150 mg/kg). Brains were rapidly removed, frozen on dry ice, and coronally sectioned. Slices containing the anterior cingulate cortex (ACC) were cryo-sectioned, yielding two 20-µm and six 100-µm sections per sample, thaw-mounted onto poly-L-lysine–coated slides.

The 20-µm sections were Nissl-stained to confirm ACC boundaries using a macaque brain atlas^78^. Tissue punches (100-µm sections) were collected on dry ice and stored at –80 °C until RNA extraction. Samples were homogenized using an M-tube (Miltenyi Biotec), and RNA was extracted with the AllPrep DNA/RNA/miRNA Universal Kit (QIAGEN, Cat# 80224).

RNA quality was verified before library construction using the Illumina® TruSeq Stranded mRNA LT Sample Prep Kit (Cat# 20020594), including mRNA enrichment, fragmentation, and reverse transcription to generate double-stranded DNA fragments (∼120–200 bp). Following end repair, adapter ligation, and PCR amplification, libraries were evaluated on an Agilent D1000 ScreenTape system (Agilent Technologies, Germany), where a single peak near 280 bp indicated optimal fragment distribution. Libraries were sequenced on an Illumina HiSeq 4000 PE100.

Reads were aligned to the *Macaca fascicularis* 5.0 genome (National Center for Biotechnology Information -NCBI; accession GCF_000364345.1) using STAR with the “quantMode GeneCounts” option. Gene-level counts were normalized in the edgeR^79^, using TMM normalization (“calcNormFactors” function). Genes with <1 count per million in at least three samples were excluded.

### Differential expression analysis

Differential expression between saline and betamethasone-treated macaques was assessed using edgeR^79^ with a false discovery rate (FDR) < 0.05. Results were visualized with EnhancedVolcano (https://bioconductor.posit.co/packages/3.19/bioc/html/EnhancedVolcano.html), highlighting genes with FDR<0.05 and |log₂ fold change| > 1.

Significantly differentially expressed genes (DEGs, FDR < 0.05) were subjected to functional enrichment analysis using MetaCore (Clarivate Analytics) and the Database for Annotation, Visualization and Integrated Discovery (DAVID, v2023q4, https://davidbioinformatics.nih.gov/)^80^, to up- and downregulated genes analyzed separately, and significance defined as FDR < 0.05.

To complement overrepresentation analyses, Gene Set Enrichment Analysis (GSEA) was performed using the fGSEA (v1.20.0, https://bioconductor.org/packages/release/bioc/html/fgsea.html), which implements a fast algorithm for the pre-ranked GSEA method ^81^. Genes were ranked by log₂ fold change between betamethasone and saline groups. Enrichment was tested against Gene Ontology Biological Process terms retrieved from Molecular Signature Database (MSigDB, v25.1.1), using 10,000 permutations. Normalized enrichment scores (NES) and FDR values were calculated, with significance defined as FDR < 0.05. Enrichment results were visualized in RStudio using ggplot2 (https://ggplot2.tidyverse.org/), ranking pathways by NES (positive and negative values indicating up- and downregulation, respectively).

#### Gene co-expression network analysis

Raw RNA-sequencing data from ACC macaques were preprocessed by removing lowly expressed genes (counts <0.9 in ≥90% of samples). Filtered data were normalized to reads per kilobase per million mappable reads [RPKM=10^9^ C/(NL)], where C is the gene’s read count, N is the total number of mapped reads, and L is the mature transcript length^82^. The normalized data were then log_2_-transformed. The preprocessed dataset (13,820 genes) was used as input for Weighted Gene Co-expression Network Analysis (WGCNA), implemented through the WGCNA R package (v.1.73)^42,83^. This approach enabled the identification of gene co-expression modules and the selection of those associated with glucocorticoid-induced transcriptional patterns.

A signed weighted network was generated by computing pairwise Pearson correlation matrix across all genes. The soft-thresholding power (β) was then selected by evaluating the scale-free topology criterion over a range of candidate β values, using the correlation matrix as input. The value β = 13 was determined using an R² cutoff of 0.90, providing the best approximation to a scale-free topology (R² = 0.9, slope = - 1.53) ^42,83^. Once the soft-threshold was defined, the correlation matrix was transformed into a signed adjacency matrix using the standard WGCNA formulation ^42,83^. The adjacency matrix was subsequently converted into a Topological Overlap Matrix (TOM) to incorporate shared network neighbours and enhance network robustness prior to module detection. Hierarchical clustering of TOM-based dissimilarity values was followed by module detection using the dynamic tree cutting (minimum module size = 30). Modules with eigengene similarity above the merging threshold (mergeCutHeight = 0.25) were merged and assigned unique color labels.

Module–trait relationships were evaluated by correlating module eigengenes (MEs) (the first principal component summarizing each module’s expression profile) with betamethasone exposure^42,51^. The correlation between each module eigengene and the experimental trait (betamethasone exposure) reflects both the magnitude and the direction of their association.

Strong eigengene–trait correlations indicate that genes within the module exhibit a coordinated expression pattern that consistently increases (or decreases) in relation to the treatment level. Modules with significant positive ME–trait associations (p < 0.05) were retained for subsequent analyses.

#### Module preservation analysis

Module preservation analysis was performed using the “*modulePreservation*” function from the WGCNA R package to assess cross-species preservation and network robustness^52^. This procedure assesses whether the co-expression module identified in the reference network (RNA-Seq data from macaques) shows comparable topological properties in independent test datasets (human postmortem cingulate cortex tissue from both healthy individuals and those with psychiatric disorders). Increased overlap and concordance across species enhance confidence in the biological relevance of the inferred networks ^50^.

The analysis quantifies preservation based on two complementary classes of statistics metrics: (i) module density, which assesses whether the overall strength of connections among genes within a module is similar in the reference and test datasets, and (ii) intramodular connectivity preservation, which evaluates whether individual genes within the module retain comparable relative connectivity within the module across datasets ^52^. Significance was assessed using a permutation-based framework (500 permutations), summarized by Zsummary and MedianRank. Zsummary > 10 indicates strong preservation, 2–10 moderate preservation, and <2 no evidence of preservation, while lower MedianRank values indicate stronger preservation.

Module preservation was assessed across three independent human post-mortem cingulate cortex datasets, each analyzed separately by diagnostic condition: (i) ACC RNA-seq from individuals with major depression, schizophrenia, bipolar disorder, and matched controls (N = 24/group)^53^; (ii) ACC tissue from controls (N = 23) and suicide cases with history of severe child abuse (N = 27)^54^; (iii) Cingulate gyrus tissue from controls (N = 21) and individuals with major depressive disorder (N = 25)^55^. All datasets were mapped to macaque Ensembl gene IDs using biomaRt^84,85^ and processed following the same filtering and normalization pipeline as the reference macaque dataset to ensure methodological consistency.

Modules that showed significant association with glucocorticoid exposure in the original macaque dataset and displayed strong preservation in human datasets were prioritized as biologically relevant for cross-species interpretation. Differences in preservation between diagnostic and control groups were quantified using ΔZsummary, defined as the difference in Zsummary values between disease and the corresponding control samples for the same module.

#### Enrichment analysis of WGCNA modules using DEGs

Enrichment of WGCNA modules for differentially expressed genes (DEGs; FDR <0.05) was assessed using gene-set overlap analysis implemented in GeneOverlap (v1.30, https://bioconductor.org/packages/GeneOverlap). Overlap between each module and DEG list was tested using Fisher’s exact test, with the background defined as all genes included in the WGCNA analysis (N = 13,820). Multiple testing correction was applied across modules, and enrichment was considered significant at FDR <0.05. Results were visualized using circular summary plots generated with Circlize^86^.

#### Module refinement by DEG-based filtering

To define biologically relevant glucocorticoid-responsive gene sets, WGCNA modules significantly associated with glucocorticoid exposure were filtered using DEG results from the same dataset. Specifically, genes belonging to each module were retained if they were differentially expressed at FDR <0.1. This filtering strategy prioritized genes exhibiting both strong co-expression and statistically robust transcriptional regulation by glucocorticoids, reducing inclusion of indirectly co-expressed but non-responsive genes.

#### Expression-based polygenic scores (ePGS)

The expression-based polygenic score (ePGS) is a biologically informed framework for analyzing genomic data that leverages gene sets defined by coordinated expression within a specific tissue^43–45,47,87,88^. In this study, ePGS were derived from human orthologs of the glucocorticoid (GC)-responsive ACC gene modules identified in macaques. Specifically, we focused on refined WGCNA modules that were positively associated with GC exposure, strongly preserved in human brain datasets, and restricted to genes differentially expressed following GC treatment (FDR < 0.1).

For each gene, the sign of the log fold change (LogFC) in response to GC exposure was incorporated into the calculation to capture the directionality of co-expression within the module. All SNPs located on these genes were retrieved using biomaRt (GRCh37.p13)^56^ and overlapped with available variants identified in ACC (BA24) from the Genotype-Tissue Expression database (GTEx, https://gtexportal.org)^89,90^ . The resulting autosomal SNP set was subjected to linkage disequilibrium clumping (r² < 0.25) to obtain independent variants. For each retained SNP, the number of effect alleles was weighted by the GTEx-estimated effect size of the SNP on gene expression. These weights were further modulated by the direction of GC-induced expression change (logFC) observed in the macaque RNA-seq data. The weighted alleles were summed across SNPs to generate individual-level ePGS values (**Figure 4A**).

Using this framework, we generated an ACC Blue-ePGS based on the refined Blue module (96 genes and 6987 SNPs). Higher ePGS values reflect stronger genetically inferred upregulation of the GC-responsive network, based on two features of the analytical framework: (i) the WGCNA modules were derived from a signed network and were positively correlated with glucocorticoid exposure, indicating coordinated transcriptional upregulation with exposure; and (ii) the directionality of glucocorticoid effects was explicitly incorporated into SNP weighting through logFC-based modulation.

To assess biological and directional specificity, ePGS were additionally constructed using the same pipeline for two comparison modules: (i) Green module ePGS/ACC derived from the refined Green module (52 genes; 2,406 SNPs), which shows positive associations with GC exposure (r = 0.77, p = 0.003) but moderate preservation in human datasets; (ii) Tan module ePGS/ACC derived from the refined tan module (23 genes; 680 SNPs), which is negatively associated with GC exposure (r = −0.87, p = 2 × 10⁻⁴). All polygenic scores were computed using PLINK 2.0^91^(www.cog-genomics.org/plink/2.0/).

#### Human genotyping data

UK Biobank blood samples were genotyped at the Affymetrix Research Services Laboratory (Santa Clara, USA). Genotyping was performed using two closely matched arrays: the bespoke BiLEVE Axiom and Affymetrix UK Biobank Axiom arrays, which share more than 95% of their marker content. In total, genotype data were generated for 487,409 participants. Axiom Array plates were processed on the Affymetrix GeneTitan® Multi-Channel Instrument. Genotypes were then called from the resulting intensities in batches of approximately 4,700 samples (around 4,800 including controls) using the Affymetrix Power Tools software and the Affymetrix Best Practices Workflow. Individuals with identical genotypes at any given SNP clustered together in a two-dimensional intensity space (one axis per targeted allele). For the interim data release, Affymetrix performed additional rounds of genotype calling with algorithms customized for the UK Biobank project, focusing on very rare variants (≤6 minor alleles per batch) and SNPs for which the generic calling algorithm did not perform optimally. After genotype calling, Affymetrix performed quality control in each batch separately to exclude SNPs with poor clustering properties. Variants failing Affymetrix QC thresholds in each batch were set to missing for all individuals within that batch. Hardy-Weinberg equilibrium was evaluated in each batch.

Affymetrix also checked sample quality (such as DNA concentration), and genotype calls were provided only for samples with sufficient DNA required metrics. For SNP-based QC metrics, only individuals with similar ancestry and the population structure were characterized by computing principal components using only UK Biobank individuals. The array design also incorporates coding variants across a wide spectrum of minor allele frequencies (MAFs), including rare variants (<1% MAF), and markers that provide good genome-wide coverage for imputation in European populations in the common (>5%) and low frequency (1–5%) MAF ranges. Further details regarding the UK Biobank genotyping procedures, quality control, and imputation pipelines were previously described ^92^. The population structure in the cohort was characterized using the fastPCA algorithm ^93^, and the first 40 principal components were included in downstream analyses to account for population stratification.

#### Phenotypic measures

Both the PHQ-9 and GAD-7 were available from two UK Biobank assessment instances. To maximize sample size and ensure consistency across phenotypic measures, symptom scores were derived from the first available assessment, using the second only when the first was unavailable.

### Patient Health Questionnaire -9 (PHQ-9)

The PHQ-9 is a self-report questionnaire administered in the UK Biobank through an online follow-up survey. It is widely used both for screening and for assessing the severity of depressive symptoms^94,95^. The questionnaire is based on DSM-IV diagnostic criteria and asks participants to report the frequency of symptoms experienced over the previous two weeks ^96^. In the UK Biobank, PHQ-9 items were obtained from the Mental Health Online follow-up (category 138), using the prompt: “*Over the last 2 weeks, how often have you been bothered by any of the following problems?*” The nine symptoms included: a) recent lack of interest or pleasure in doing things (#20514, #29002, anhedonia); b) recent feelings of depression (#20510, #29003, depressed mood); c)trouble falling or staying asleep, or sleeping too much (#20517, #29004, sleep problems); d) recent feelings of tiredness or low energy (#20519, #29005, tiredness); e) recent poor appetite or overeating (#20511, #29006, changes in appetite); f) recent feelings of inadequacy (#20507, #29007, feelings of inadequacy); g) recent trouble concentrating on things (#20508, #29008, concentration problems); h) recent changes in speed/amount of moving or speaking (#20518, #29009, psychomotor changes); and i) recent thoughts of suicide or self-harm (#20513, #29010, suicidality). Responses to these nine items were summed to generate a total PHQ-9 score, with scores ≥10 commonly interpreted as indicative of probable depression.

### General Anxiety Disorder-7 (GAD-7)

The same Mental Health Online follow-up (category 138) included the items comprising the GAD-7, a validated self-report measure of generalized anxiety symptom severity^97^. As with the PHQ-9, participants were asked: “*Over the last 2 weeks, how often have you been bothered by any of the following problems?*” The seven anxiety-related items were: a) Feeling nervous, anxious or on edge (#20506, #29058); 2) not being able to stop or control worrying (#20509, #29059); c) worrying too much about different things (#20520, #29060); d) trouble relaxing (#20515, #29061); e) being so restless that it is hard to sit still (#20516, #29062); f) becoming easily annoyed or irritable (#20505, #29063); g) feeling afraid as if something awful might happen (#20512, #29064) ^98^. Standard thresholds classify scores of 5, 10, and 15 as reflecting mild, moderate, and severe anxiety symptoms, respectively ^97^.

#### Adversity scores

Early-life adversity was assessed using a cumulative score derived from the Childhood Trauma Questionnaire (CTQ) in the UK Biobank^99,100^ (item-level details in **Table S8)**. CTQ data were available from two assessment instances. To maximize sample size, CTQ scores were derived from the first available assessment, using the second only when the first was unavailable. The CTQ-based measure comprised five domains reflecting emotional neglect, physical neglect, and experiences of emotional, physical, and sexual abuse early in life. Responses across five items were aggregated to generate a continuous early-life adversity score.

Adult adversity was quantified using a cumulative score based on five self-reported adverse experiences occurring since age 16, aggregated into a continuous adult trauma score (**Table S8**).

A combined adversity score was constructed by summing early-life and adult adversity scores to capture lifetime exposure. This composite measure reflects total adversity exposure level while preserving contributions from both early-life and adult exposures. All adversity scores were analyzed as continuous variables, with higher values indicating greater adversity exposure. Participants with missing data for a given adversity measure were excluded only from analyses involving that specific score.

#### Developmental and adult ACC gene expression patterns

Normalized gene expression data from the BrainSpan Atlas of the Developing Human Brain (http://www.brainspan.org)^90^ were used to examine developmental changes in expression patterns of the ACC GC-responsive module (refined Blue module ePGS/ACC). Human ortholog genes from the refined Blue-ePGS were matched by Ensembl ID in BrainSpan, retaining genes present across samples (95/96 genes). Analyses were restricted to anterior cingulate (medial prefrontal) cortex samples. Two developmental stages were defined based on metadata: childhood (1–13 years; N = 7) and adulthood (19–37 years; N = 6).

Expression matrices for each group were constructed after removing genes with zero variance. For each developmental group, we computed pairwise gene–gene Pearson correlations and generated clustered heatmaps using the heatmaply^101^.

To quantify developmental differences in co-expression structure, unique correlation values were extracted (upper triangle of the correlation matrix), computed absolute correlations (|r|), and the proportion of strongly co-expressed gene pairs (|r| > 0.5). Developmental differences were assessed using the Wilcoxon rank-sum test (|r| distributions), Fisher’s exact test for enrichment of high-correlation pairs (|r| > 0.5), and permutation testing (10,000 iterations) assessing whether the observed difference in mean |r| could arise under random relabeling of age groups. Distributions were visualized using ggplot2 (https://ggplot2.tidyverse.org/).

To assess co-expression patterns of the refined Blue module ePGS/ACC in the adult clinical context, a complementary analysis was performed using post-mortem ACC gene expression data^54^, corresponding to the same cohort used in the module preservation analysis. Expression data for the Blue module ePGS/ACC human orthologs were extracted (68/96 genes) and analyzed separately in control individuals (N=18) and suicide cases (N=24). Pairwise gene-gene Pearson correlation matrices were computed within each group, and clustered heatmaps were generated using identical parameters. As above, unique correlation coefficients were extracted from the upper triangle of each matrix, absolute correlations (|r|) were calculated, and the proportion of strongly co-expressed gene pairs (|r| > 0.5) was quantified for each group.

These metrics were used to compare global co-expression strength between control and suicide samples.

#### Functional characterization of the glucocorticoid-responsive module

Functional enrichment analyses of Blue module ePGS/ACC genes were performed using DAVID (v2023q4, https://davidbioinformatics.nih.gov/)^80^ , testing Gene Ontology biological processes (BP), cellular components (CC), and molecular functions (MF) terms. Human ortholog genes included in the ePGS calculation were used as input (96 genes). Enriched terms were ranked by FDR, and the top 10 terms (FDR < 0.05) were selected for interpretation. Visualizations were generated using ggplot2 (https://ggplot2.tidyverse.org). The full set of preprocessed genes (N = 13,820) was used as the background universe for all enrichment analyses.

Network topology was visualized in Cytoscape®^102^. The network was generated by importing the 96 human orthologous genes into the GeneMANIA plugin, retaining only co-expression interactions. All additional genes automatically added by GeneMANIA were removed to restrict analyses to the original GC-responsive gene set. Network centrality metrics (degree and betweenness) were computed using the CentiScaPe plugin to identify highly connected hub genes.

Genome-wide association study (GWAS) enrichment analyses were performed using MAGMA v1.10 ^103^. GWAS summary statistics were first mapped to genes using the FUMA platform^104^, including studies of (i) anxiety symptoms^105^, (ii) depressive symptoms^106^, (iii) major depressive disorder (MDD) case–control status^107^, and (iv) a multi-trait meta-analysis^108^.

MAGMA-based gene annotation was applied to derive gene-level association statistics. SNPs were assigned to genes using a 0 kb window based on the GRCh38 genome build, and gene-level association statistics were exported in MAGMA-compatible format.

Gene property analyses were subsequently performed using MAGMA to test whether gene-level GWAS associations were related to intramodular connectivity within Blue module ePGS/ACC. Specifically, module membership (kME) values from the refined ACC Blue module were used as a continuous gene-level covariate, with gene-level GWAS Z-scores regressed on kME using MAGMA’s linear regression framework. One-sided tests (positive direction) were used to assess whether higher connectivity was associated with stronger GWAS signals, with statistical significance evaluated using MAGMA’s permutation-based approach.

Gene set overlap analyses were performed to compare the ACC Blue module gene network with GC-responsive gene networks previously identified in the anterior and posterior hippocampal dentate gyrus of the same female macaques ^47^. Overlap was assessed using the direct intersection of gene lists, and results are presented descriptively without statistical testing.

## STATISTICAL ANALYSIS

All analyses were conducted in RStudio Workbench. Statistical significance was set at α < 0.05. Linear regression models were fitted to test main and interaction effects between ePGS (continuous variable) and adversity measures on PHQ-9 and GAD-7 outcomes. All continuous variables were standardized before analyses. All models were adjusted for age, assessment center, genotype array, and the first 40 genetic principal components to account for population stratification within the UK Biobank cohort. Descriptive statistics summarized baseline characteristics by refined blue ePGS groups (**Table 1**). Group comparisons were performed using a two-sided Student’s t-test for continuous variables and Pearson’s chi-squared test for categorical variables. Baseline characteristics were broadly comparable across ePGS groups.

To evaluate the biological specificity of the observed interactions, ePGS were additionally constructed for two other WGCNA modules using the same analytical pipeline applied to the Blue module ePGS/ACC, including DEG filtering at FDR < 0.1. A refined Green module ePGS/ACC was derived from a module positively associated with GC exposure (52 genes; 2,406 SNPs) but moderately preserved in human brain datasets. As a directional control, a Tan module ePGS/ACC was constructed from a module negatively associated with GC exposure (23 genes; 680 SNPs). Both comparison ePGS were tested using identical model specifications, covariates, environmental exposures, and outcome measures as in the primary analyses.

## Notes

### Competing Interest Statement

The authors have declared no competing interest.

