## Supplemental Material for "A glucocorticoid-responsive polygenic signature in the anterior cingulate cortex moderates the association of early-life adversity and vulnerability for depression"

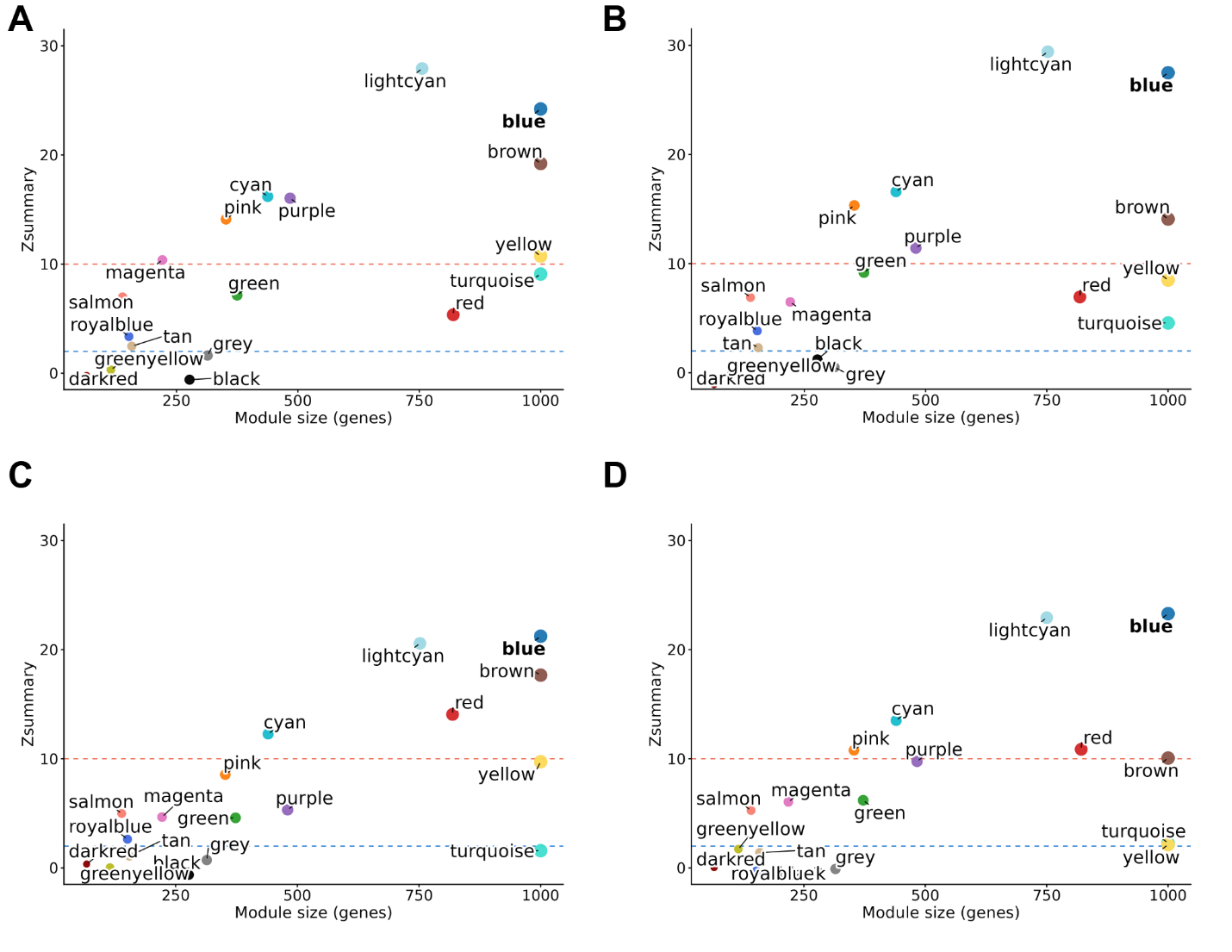

**Figure S1. Z-summary statistics for WGCNA module preservation across psychiatric conditions (related to Figure 3).** Z-summary values are shown as a function of module size (number of genes) to assess module preservation relative to the dataset from Ramaker et al. 2017 [S1]. Panels correspond to **(A)** controls, **(B)** major depressive disorder, **(C)** schizophrenia, and **(D)** bipolar disorder. Dashed horizontal lines indicate conventional thresholds for moderate (Z-summary > 2, blue) and strong (Z-summary > 10, red) module preservation.

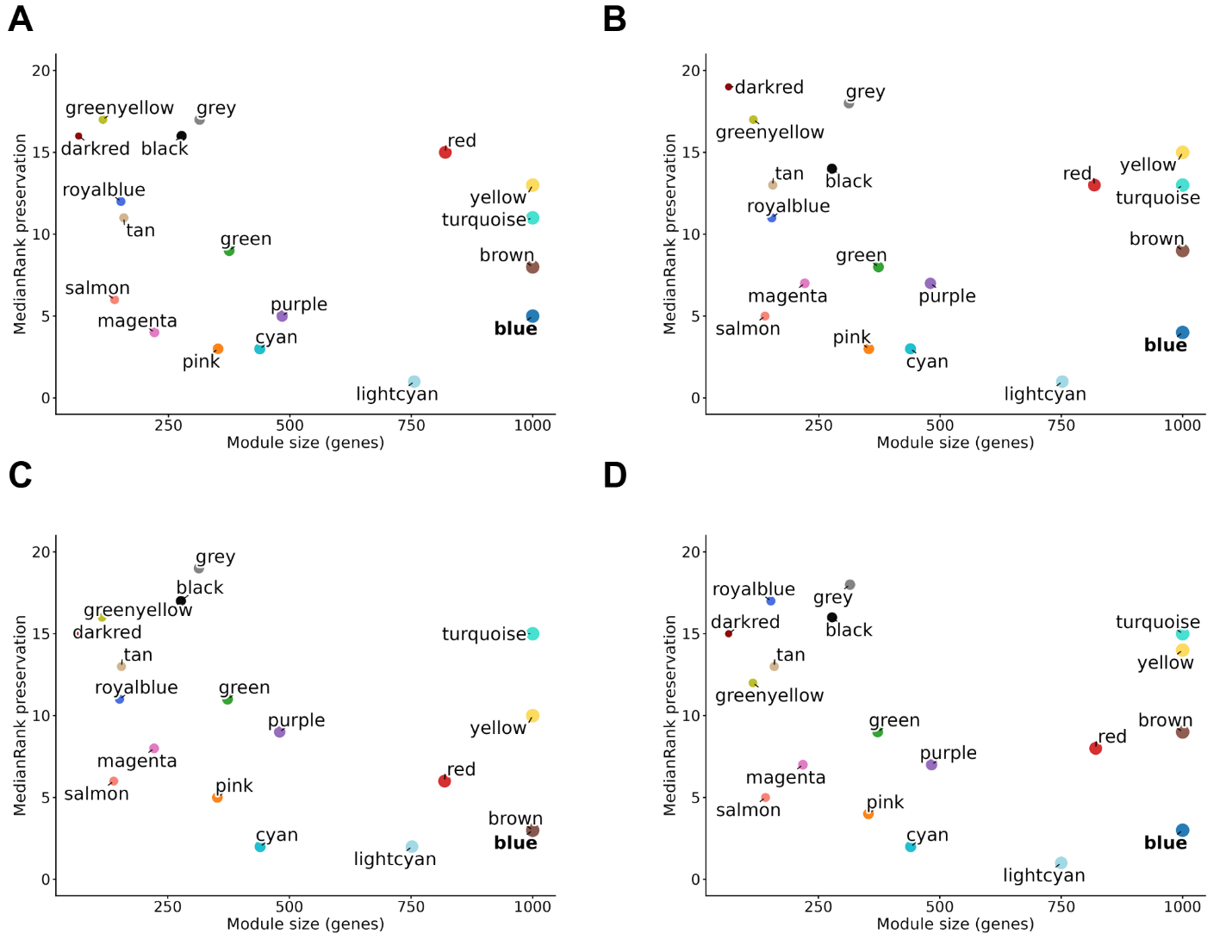

**Figure S2. MedianRank statistics for WGCNA module preservation across psychiatric conditions (related to Figure 3).** MedianRank values are shown as a function of module size (number of genes) to assess module preservation relative to the dataset from Ramaker et al. 2017 [S1]. Panels correspond to **(A)** controls, **(B)** major depressive disorder, **(C)** schizophrenia, and **(D)** bipolar disorder. Lower MedianRank values indicate stronger module preservation.

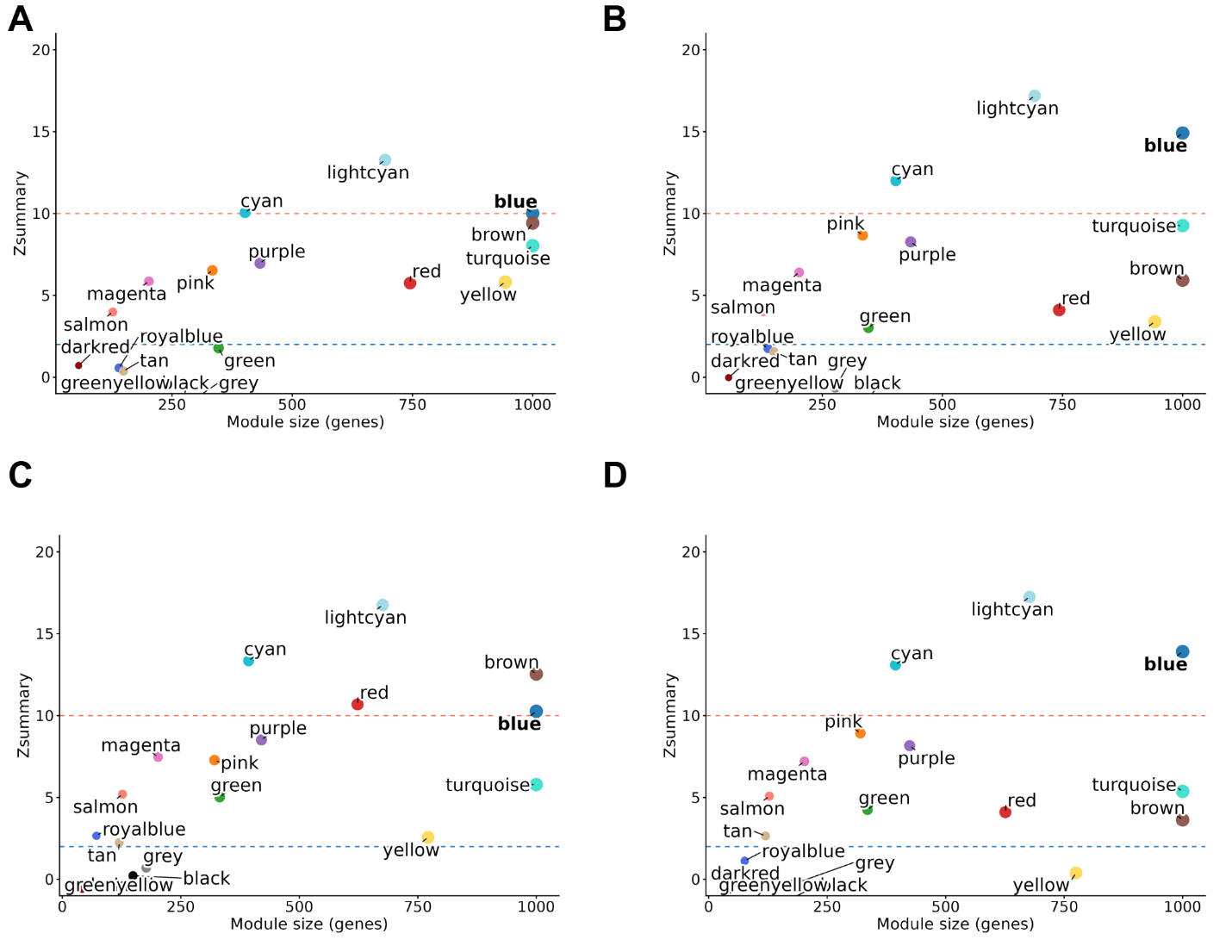

**Figure S3. Z-summary statistics for WGCNA module preservation across psychiatric conditions (related to Figure 3).** Z-summary values are shown as a function of module size (number of genes) to assess module preservation relative to the dataset to independent datasets from Lutz et al. 2017 [S2] and Labonté et al. 2017 [S3]. Panels correspond to **(A)** controls (Lutz), **(B)** suicide cases (Lutz), **(C)** controls (Labonté), and **(D)** depression (Labonté). Dashed horizontal lines indicate conventional thresholds for moderate (Z-summary > 2, blue) and strong (Z-summary > 10, red) module preservation.

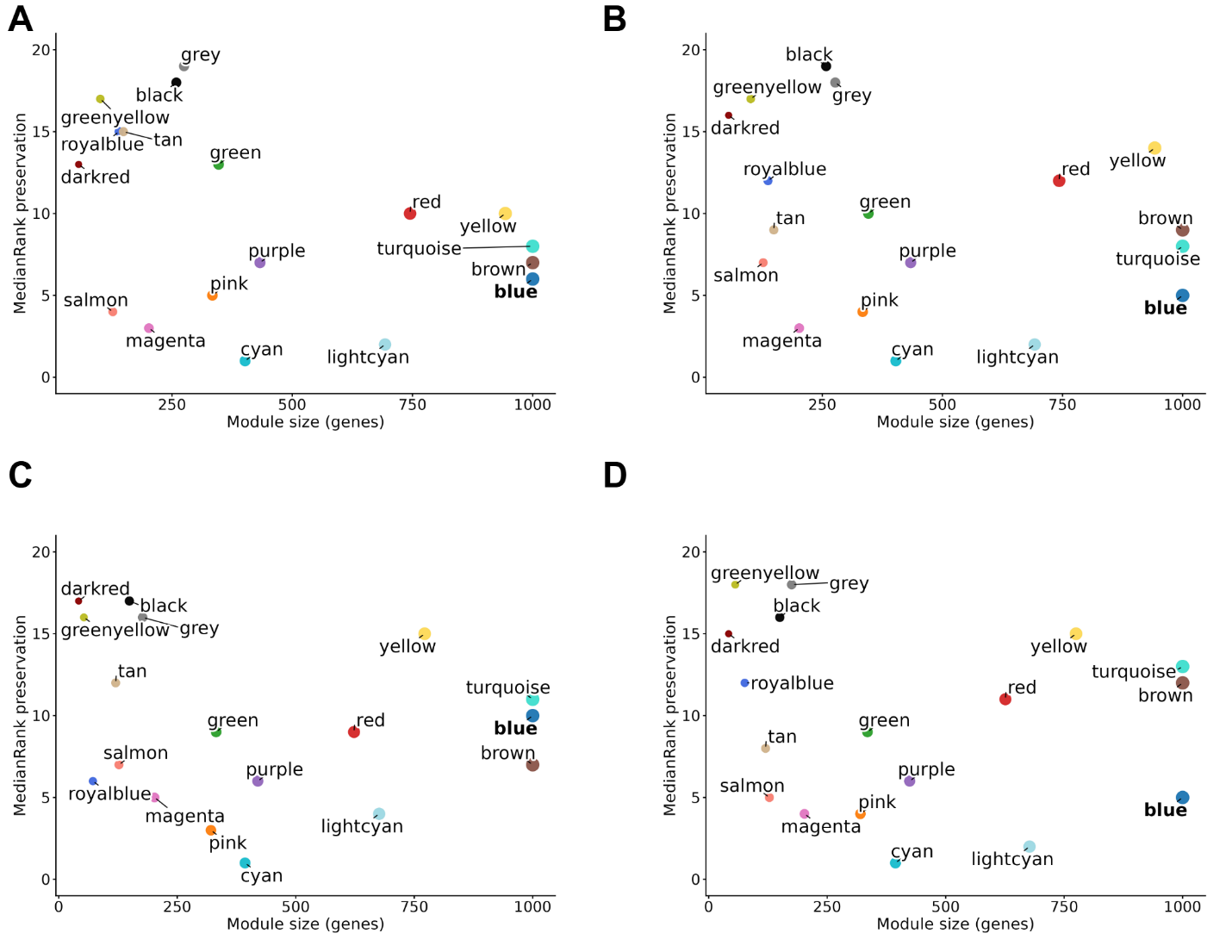

**Figure S4. MedianRank statistics for WGCNA module preservation across psychiatric conditions (related to Figure 3).** MedianRank values are shown as a function of module size (number of genes) to assess module preservation relative to the dataset from Lutz et al. 2017 [S2] and Labonté et al. 2017 [S3]. Panels correspond to **(A)** controls (Lutz), **(B)** suicide cases (Lutz), **(C)** controls (Labonté), and **(D)** depression (Labonté). Lower MedianRank values indicate stronger module preservation.

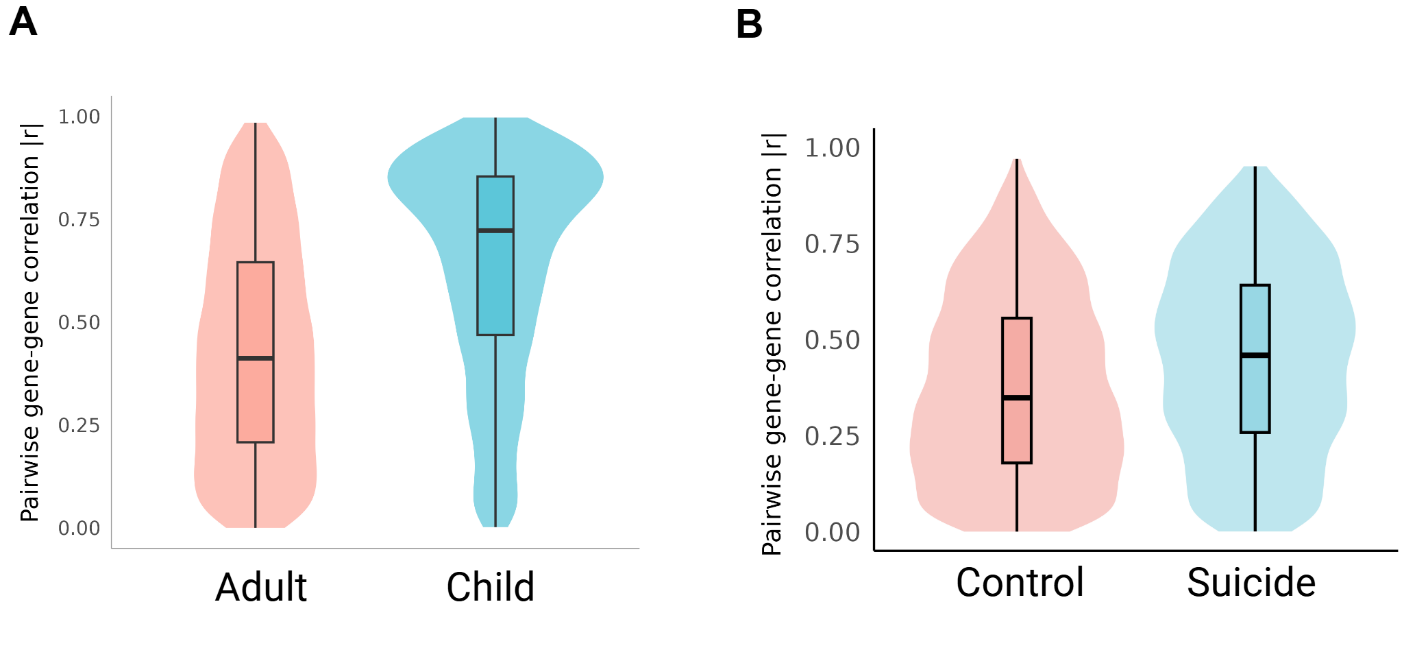

**Figure S5. Distribution of pairwise gene–gene expression correlations among Blue module ePGS/ACC genes across developmental stage and diagnostic status (related to Figure 6).** Violin plots show the distribution of pairwise gene–gene expression correlation coefficients (r) among genes in the refined blue-ePGS network. **(A)** Comparison between adult and childhood samples from BrainSpan. **(B)** Comparison between control individuals and suicide cases from Lutz et al. 2017 [S2]. Boxplots indicate the median and interquartile range.

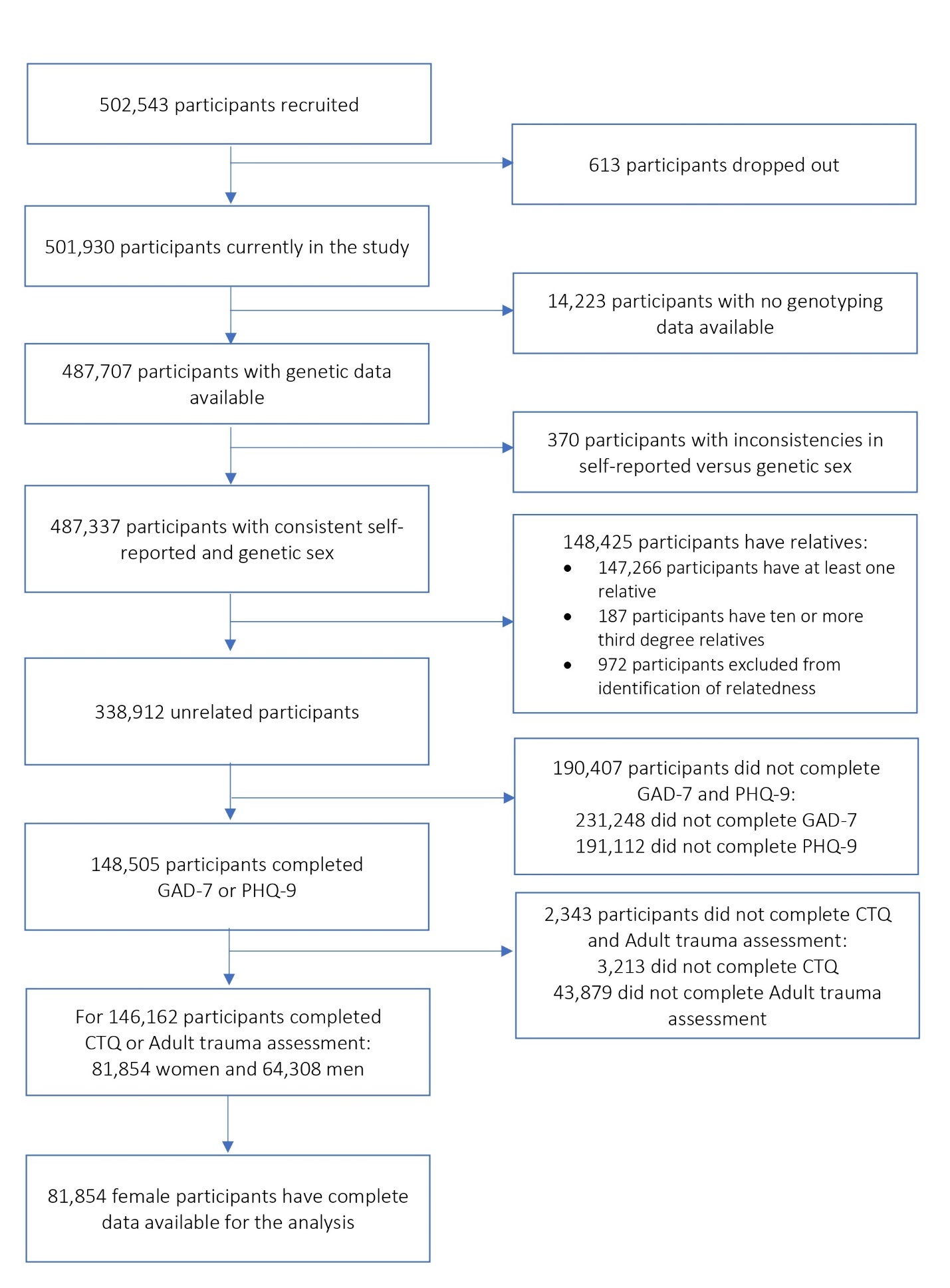

**Figure S6. Flowchart of participant selection in the UK Biobank (related to STAR Methods).** Sequential inclusion and exclusion criteria applied to derive the final analytic sample of females with complete genetic, adversity, and depressive symptom data.

**Table S1. Differentially expressed genes in the macaque anterior cingulate cortex (ACC) following betamethasone versus saline treatment (FDR < 0.05).** Related to Figure 1.

| Gene | LogFC | P-value | FDR |
| --- | --- | --- | --- |
| *PTK2B* | 0.68144506 | 5.78E-16 | 9.21E-12 |
| *LCN9* | 3.694155918 | 1.44E-13 | 1.14E-09 |
| *MSMO1* | -0.694075668 | 2.59E-12 | 1.38E-08 |
| *ZBTB16* | 0.642699238 | 9.65E-12 | 3.84E-08 |
| *SULT1E1* | 2.25420287 | 3.80E-11 | 1.21E-07 |
| *SAMD14* | 0.754629041 | 8.32E-11 | 2.21E-07 |
| *APOD* | 0.852998703 | 2.76E-10 | 6.28E-07 |
| *CD74* | -1.061746416 | 3.98E-10 | 7.92E-07 |
| *LGI3* | 0.546980407 | 8.10E-10 | 1.43E-06 |
| *CNDP1* | -1.660956293 | 9.70E-10 | 1.54E-06 |
| *LTB* | 1.919515169 | 1.22E-09 | 1.75E-06 |
| *PCSK1* | 0.71744775 | 1.32E-09 | 1.75E-06 |
| *KCNS1* | 0.756982033 | 1.59E-09 | 1.95E-06 |
| *LRRC17* | -1.429652242 | 1.76E-09 | 2.00E-06 |
| *LOC102144039* | -1.277335728 | 2.43E-09 | 2.58E-06 |
| *DYDC2* | -0.897277033 | 6.04E-09 | 6.01E-06 |
| *RHBDL3* | 0.820259429 | 2.13E-08 | 1.99E-05 |
| *IFI44* | -1.247804998 | 3.17E-08 | 2.81E-05 |
| *C1QTNF3* | -1.245173353 | 1.66E-07 | 0.000139 |
| *SFRP4* | -1.263311823 | 1.97E-07 | 0.000157 |
| *CARTPT* | 1.387340637 | 3.60E-07 | 0.000273 |
| *OAS2* | -1.086953362 | 4.51E-07 | 0.000325 |
| *IGFBP6* | 0.767021163 | 4.87E-07 | 0.000325 |
| *CCDC152* | -0.484212904 | 4.90E-07 | 0.000325 |
| *LOC102121035* | -2.40180876 | 5.63E-07 | 0.000356 |
| *LOC102136049* | -2.602088026 | 5.81E-07 | 0.000356 |
| *FKBP5* | 0.952519321 | 6.57E-07 | 0.000377 |
| *PCP2* | -3.950536614 | 6.64E-07 | 0.000377 |
| *TUBA1A* | -0.487927529 | 9.17E-07 | 0.000502 |
| *HECW1* | 0.527184417 | 9.46E-07 | 0.000502 |
| *DUSP4* | 0.633338773 | 1.08E-06 | 0.000556 |
| *ABCG2* | -0.724499293 | 1.12E-06 | 0.000559 |
| *KCNB1* | 0.638514592 | 1.33E-06 | 0.000643 |
| *GFOD1* | 1.058816131 | 1.41E-06 | 0.000659 |
| *CD163* | 0.888131603 | 1.50E-06 | 0.000681 |
| *LOC102119202* | 0.453736247 | 1.71E-06 | 0.000755 |
| *ANKRD34C* | 0.641867087 | 2.28E-06 | 0.000958 |
| *LMO2* | -0.6283679 | 2.29E-06 | 0.000958 |
| *CCDC129* | 1.65929319 | 2.49E-06 | 0.001015 |
| *KCNA2* | 0.493327623 | 2.59E-06 | 0.001029 |
| *PLCH1* | 0.430807545 | 2.93E-06 | 0.001137 |
| *CXCL12* | -0.75168548 | 3.00E-06 | 0.001137 |
| *CALB2* | -0.526611402 | 3.31E-06 | 0.001225 |
| *SIK2* | 0.481593325 | 3.46E-06 | 0.001251 |
| *ACAT2* | -0.521038339 | 3.77E-06 | 0.001333 |
| *CIT* | 0.361567141 | 4.23E-06 | 0.001462 |
| *CRLF1* | 1.224160569 | 4.65E-06 | 0.001573 |
| *PDZD2* | 0.51647979 | 5.02E-06 | 0.001664 |
| *GGACT* | 0.888574406 | 5.17E-06 | 0.001679 |
| *LOC102128633* | 1.34180187 | 5.75E-06 | 0.001831 |
| *PI16* | -1.228822339 | 6.85E-06 | 0.002136 |
| *BCL6* | 0.431597114 | 7.84E-06 | 0.002398 |
| *SLC5A3* | 0.545459898 | 8.30E-06 | 0.002493 |
| *RORB* | 0.816382354 | 8.79E-06 | 0.002591 |
| *LOC102142617* | -0.804067907 | 8.97E-06 | 0.002595 |
| *CX3CR1* | -1.168833522 | 9.66E-06 | 0.002745 |
| *CTXN3* | 1.000811153 | 1.05E-05 | 0.002929 |
| *TNFSF10* | -0.953999305 | 1.12E-05 | 0.003071 |
| *HSD11B1* | -1.411794075 | 1.14E-05 | 0.003082 |
| *LOC107129898* | 1.298822714 | 1.20E-05 | 0.003188 |
| *IGSF1* | -0.786541482 | 1.29E-05 | 0.003361 |
| *CD52* | -0.922294557 | 1.38E-05 | 0.003517 |
| *LOC102137548* | 1.053576051 | 1.39E-05 | 0.003517 |
| *PRKCA* | 0.770359515 | 1.56E-05 | 0.003888 |
| *STMN1* | -0.450900216 | 1.74E-05 | 0.004263 |
| *SPEN* | 0.415928132 | 1.80E-05 | 0.004348 |
| *RASGRF1* | 0.435912246 | 1.91E-05 | 0.004425 |
| *AIF1* | -0.907261159 | 1.91E-05 | 0.004425 |
| *BTBD3* | 0.438927497 | 1.92E-05 | 0.004425 |
| *PODN* | 0.814523618 | 2.07E-05 | 0.004695 |
| *LOC107126405* | 1.16010795 | 2.16E-05 | 0.004837 |
| *NMRAL1* | -0.643319337 | 2.66E-05 | 0.005848 |
| *CCDC3* | 0.647913987 | 2.68E-05 | 0.005848 |
| *LOC102123433* | 1.691123699 | 3.68E-05 | 0.007921 |
| *UNC13A* | 0.362514445 | 4.01E-05 | 0.008511 |
| *LOC102131151* | -0.48439227 | 4.08E-05 | 0.008525 |
| *TNS1* | 0.473480533 | 4.12E-05 | 0.008525 |
| *RAPGEF5* | 0.415789198 | 4.37E-05 | 0.008801 |
| *KLF9* | 0.703284388 | 4.37E-05 | 0.008801 |
| *SQLE* | -0.429462842 | 4.47E-05 | 0.008892 |
| *LOC102145743* | -0.683363248 | 4.80E-05 | 0.009307 |
| *CAMK1G* | 0.379855543 | 4.82E-05 | 0.009307 |
| *TNR* | 0.743320343 | 4.85E-05 | 0.009307 |
| *ACOT2* | -0.596848946 | 5.11E-05 | 0.00955 |
| *ST6GAL2* | -0.458141247 | 5.12E-05 | 0.00955 |
| *KCNS2* | 0.423242415 | 5.16E-05 | 0.00955 |
| *AIF1L* | -0.587928912 | 5.36E-05 | 0.009773 |
| *SORL1* | 0.351310809 | 5.40E-05 | 0.009773 |
| *IDI1* | -0.397443095 | 5.55E-05 | 0.009932 |
| *MYBPH* | 1.038112972 | 5.95E-05 | 0.010485 |
| *P2RY12* | -0.961025825 | 6.00E-05 | 0.010485 |
| *MATN3* | 1.446297763 | 6.40E-05 | 0.011077 |
| *B2M* | -0.522701064 | 6.74E-05 | 0.01153 |
| *LOC107130744* | -0.547781592 | 7.22E-05 | 0.012217 |
| *SPRY4* | 0.447059952 | 7.85E-05 | 0.012961 |
| *PLAC8* | -1.366198606 | 7.87E-05 | 0.012961 |
| *KLHL3* | 0.410681659 | 7.90E-05 | 0.012961 |
| *RBM3* | -0.458002779 | 8.07E-05 | 0.013104 |
| *WIPF3* | 0.366701456 | 8.55E-05 | 0.013667 |
| *VPS53* | 0.514167595 | 8.61E-05 | 0.013667 |
| *CTHRC1* | -0.948698772 | 8.67E-05 | 0.013667 |
| *UBD* | -2.065165123 | 9.07E-05 | 0.014074 |
| *SLC36A1* | 0.558245141 | 9.11E-05 | 0.014074 |
| *HSD17B6* | -0.559051647 | 9.45E-05 | 0.014452 |
| *ZBTB40* | 0.382606989 | 9.54E-05 | 0.014452 |
| *SLC35E2B* | 0.452669642 | 9.65E-05 | 0.014494 |
| *FAM212B* | 0.541043188 | 9.87E-05 | 0.014674 |
| *SLC37A2* | -0.676536032 | 0.000106 | 0.015567 |
| *LOC102137553* | 0.814241663 | 0.000107 | 0.015567 |
| *PROS1* | 0.746176398 | 0.000113 | 0.01623 |
| *LOC107128414* | 1.055751087 | 0.000113 | 0.01623 |
| *IGFBP5* | 1.178982226 | 0.000119 | 0.016822 |
| *CCNJL* | -0.690626456 | 0.000119 | 0.016822 |
| *KCND3* | 0.900876813 | 0.000124 | 0.017114 |
| *MGAT5* | 0.713298107 | 0.000124 | 0.017114 |
| *LOC102136468* | -1.253445452 | 0.000125 | 0.017114 |
| *ARPP19* | -0.347225481 | 0.000139 | 0.018928 |
| *MAS1* | 0.943718299 | 0.000142 | 0.019175 |
| *FOSL2* | 0.349610138 | 0.000144 | 0.019297 |
| *STX1B* | 0.377326384 | 0.000154 | 0.020432 |
| *LOC102128162* | -1.193628307 | 0.000159 | 0.020937 |
| *LOC102141262* | -1.458366616 | 0.000166 | 0.021441 |
| *ARHGAP25* | -0.904328095 | 0.000166 | 0.021441 |
| *LOC102132369* | -1.012438554 | 0.000176 | 0.02252 |
| *FGFBP3* | -0.628878445 | 0.000177 | 0.02252 |
| *SYNJ2* | 0.356479029 | 0.000178 | 0.02252 |
| *LOC102134040* | 0.360268832 | 0.000182 | 0.022779 |
| *MKLN1* | 0.474429202 | 0.000183 | 0.022779 |
| *LOC102142698* | 0.406822314 | 0.000192 | 0.023571 |
| *LOC102141176* | -1.491902794 | 0.000193 | 0.023571 |
| *EDN3* | -0.890614535 | 0.000195 | 0.023604 |
| *UCP2* | -0.714006166 | 0.000196 | 0.023604 |
| *SOGA3* | 0.736421826 | 0.000199 | 0.02386 |
| *ALDH4A1* | 0.55716984 | 0.000204 | 0.02424 |
| *EMILIN3* | 0.815542471 | 0.000211 | 0.024593 |
| *BSN* | 0.314780799 | 0.000212 | 0.024593 |
| *FABP3* | -0.38683574 | 0.000213 | 0.024593 |
| *RAMP2* | -0.557550557 | 0.000213 | 0.024593 |
| *TUSC3* | -0.317836209 | 0.000216 | 0.024616 |
| *PLEKHM3* | 0.809257774 | 0.000217 | 0.024616 |
| *SHC3* | 0.345727512 | 0.000219 | 0.024713 |
| *KDM5C* | 0.352919599 | 0.000224 | 0.025104 |
| *NAV1* | 0.371298475 | 0.000226 | 0.025109 |
| *TTR* | -1.286716858 | 0.00023 | 0.025228 |
| *ANKRD33B* | 0.501844493 | 0.00023 | 0.025228 |
| *SLC45A4* | 0.511468012 | 0.000231 | 0.025228 |
| *ETV5* | 0.517544099 | 0.000238 | 0.025742 |
| *PLP1* | -0.735072849 | 0.000241 | 0.025742 |
| *TDRD9* | 0.883766977 | 0.000241 | 0.025742 |
| *ESD* | -0.389139651 | 0.000243 | 0.025781 |
| *LOC102133457* | -1.048449207 | 0.000245 | 0.02586 |
| *FOXF2* | -0.737992619 | 0.00025 | 0.026041 |
| *PEA15* | -0.316775431 | 0.000251 | 0.026041 |
| *C14H11orf87* | 0.571555903 | 0.000253 | 0.026041 |
| *CETN2* | -0.367987272 | 0.000254 | 0.026041 |
| *FLVCR2* | -0.984227442 | 0.000261 | 0.026661 |
| *HMGCS1* | -0.376637732 | 0.000268 | 0.026921 |
| *HOMER1* | 0.423344469 | 0.000269 | 0.026921 |
| *ADCY1* | 0.611036299 | 0.000269 | 0.026921 |
| *IPCEF1* | 0.385224341 | 0.000272 | 0.026921 |
| *FRRS1L* | 0.752516371 | 0.000273 | 0.026921 |
| *SIDT2* | 0.360573737 | 0.000274 | 0.026921 |
| *CHRNB2* | 0.369571033 | 0.000279 | 0.027255 |
| *IFI44L* | -1.315358831 | 0.000298 | 0.028882 |
| *BEX4* | -0.323979526 | 0.000305 | 0.029255 |
| *LRRC8B* | 0.456275742 | 0.000306 | 0.029255 |
| *JCHAIN* | -3.203760535 | 0.000307 | 0.029255 |
| *C1QC* | 0.875384263 | 0.000312 | 0.029519 |
| *TTBK2* | 0.578426567 | 0.000319 | 0.030032 |
| *NENF* | -0.438003444 | 0.000322 | 0.030167 |
| *STRBP* | 0.304275804 | 0.000328 | 0.030357 |
| *TET3* | 0.352274959 | 0.000328 | 0.030357 |
| *VEGFA* | 0.412499311 | 0.00033 | 0.030357 |
| *TCP11L1* | 0.626468336 | 0.000333 | 0.030447 |
| *OCRL* | 0.412455394 | 0.000336 | 0.030558 |
| *USH1C* | 0.626865029 | 0.000344 | 0.031071 |
| *FRY* | 0.336857486 | 0.000359 | 0.031826 |
| *KIAA1549* | 0.501816661 | 0.00036 | 0.031826 |
| *MED28* | 0.567423006 | 0.000363 | 0.031826 |
| *ABCB1* | -0.770167673 | 0.000365 | 0.031826 |
| *C5H4orf50* | 0.647902949 | 0.000366 | 0.031826 |
| *IGSF9B* | 0.475686392 | 0.000366 | 0.031826 |
| *FAM162A* | -0.372672268 | 0.000367 | 0.031826 |
| *SLC17A8* | -1.467799902 | 0.000369 | 0.031826 |
| *FTH1* | -0.360324668 | 0.000371 | 0.031826 |
| *CRYM* | -0.420371187 | 0.000372 | 0.031826 |
| *KYNU* | -0.80553118 | 0.000376 | 0.031987 |
| *TSSK2* | 1.002996728 | 0.000379 | 0.032075 |
| *LOC102115534* | 0.282065815 | 0.000382 | 0.032143 |
| *BTG2* | -0.397743521 | 0.000384 | 0.032145 |
| *KCNH1* | 0.40114235 | 0.000386 | 0.032145 |
| *TNMD* | -1.395017856 | 0.000399 | 0.03311 |
| *FDPS* | -0.364205769 | 0.000407 | 0.033596 |
| *TMX4* | 0.681992183 | 0.00041 | 0.0336 |
| *LUZP1* | 0.356424125 | 0.000414 | 0.033784 |
| *CYSLTR1* | -1.033363798 | 0.000428 | 0.034728 |
| *FLI1* | -0.713388524 | 0.000437 | 0.034993 |
| *PLXNA4* | 0.509863678 | 0.000437 | 0.034993 |
| *ZNF189* | 0.398236699 | 0.000438 | 0.034993 |
| *MARCKS* | -0.293139286 | 0.000446 | 0.035408 |
| *LRRK1* | 0.581954584 | 0.000448 | 0.035408 |
| *CAMK1D* | 0.433810883 | 0.000449 | 0.035408 |
| *TEK* | -0.665158383 | 0.000453 | 0.035519 |
| *FBXO4* | -0.659980568 | 0.00046 | 0.035913 |
| *CLSTN2* | 0.614725722 | 0.000482 | 0.037443 |
| *VSIG10L* | 0.909411676 | 0.000488 | 0.037731 |
| *P2RY13* | -0.703836668 | 0.000513 | 0.039448 |
| *PAPLN* | 0.607008585 | 0.00052 | 0.039785 |
| *POSTN* | -0.906122824 | 0.000525 | 0.039938 |
| *LOC107127758* | -0.835420964 | 0.000528 | 0.03995 |
| *NUDT10* | -0.385524815 | 0.00053 | 0.03995 |
| *MAP9* | 0.610743245 | 0.000553 | 0.041359 |
| *LOC107130982* | -0.803303563 | 0.000554 | 0.041359 |
| *NCKIPSD* | 0.388179516 | 0.000569 | 0.042271 |
| *LOC102141871* | -1.840279549 | 0.000571 | 0.042271 |
| *CAMK2A* | 0.298189541 | 0.000588 | 0.043159 |
| *KCNAB2* | 0.45995396 | 0.000589 | 0.043159 |
| *LOC107126719* | -1.486318737 | 0.000597 | 0.043614 |
| *SV2C* | 0.98298417 | 0.000605 | 0.043728 |
| *ATCAY* | 0.351851274 | 0.000607 | 0.043728 |
| *PBK* | -1.030278144 | 0.000607 | 0.043728 |
| *KIAA0513* | 0.445347286 | 0.00061 | 0.043728 |
| *TXN* | -0.369932553 | 0.000617 | 0.043952 |
| *RCN1* | 0.381940159 | 0.00062 | 0.043952 |
| *C11H12orf42* | 0.727664732 | 0.000626 | 0.043952 |
| *KDM7A* | 0.495017622 | 0.000626 | 0.043952 |
| *MAPK13* | 0.303522543 | 0.000627 | 0.043952 |
| *SORT1* | 0.496340093 | 0.000633 | 0.044214 |
| *UGGT1* | 0.389595064 | 0.000646 | 0.044761 |
| *MAPK4* | 0.282473714 | 0.000647 | 0.044761 |
| *TUBB* | -0.290832355 | 0.00066 | 0.045476 |
| *MFSD8* | -0.379693317 | 0.000665 | 0.045593 |
| *HECTD4* | 0.333713862 | 0.000674 | 0.04603 |
| *SLAIN1* | -0.361129288 | 0.000685 | 0.046569 |
| *HR* | 0.585468727 | 0.000696 | 0.047116 |
| *IFI6* | -0.442136324 | 0.000703 | 0.047303 |
| *PLXDC1* | -0.416380521 | 0.000704 | 0.047303 |
| *EVI2A* | -0.715664213 | 0.000711 | 0.047381 |
| *SLC38A1* | 0.641571008 | 0.000712 | 0.047381 |
| *USP37* | 0.760398057 | 0.000722 | 0.047758 |
| *HTT* | 0.337835346 | 0.000726 | 0.047758 |
| *NXPH3* | 0.555904611 | 0.000727 | 0.047758 |
| *CFB* | 0.93138727 | 0.000729 | 0.047758 |
| *TM4SF18* | -0.783060944 | 0.000759 | 0.04929 |
| *ITPR1* | 0.365385034 | 0.000759 | 0.04929 |
| *FBXL18* | 0.675179118 | 0.000764 | 0.049393 |

**Table S2. List of human ortholog genes comprising the refined Blue, Green, and Tan modules ePGS/ACC.** Related to Figures 3 and 4A.

| Blue module ePGS | Green module ePGS | Tan module ePGS |
| --- | --- | --- |
| *ADCY1* | *AGO2* | *ANGPT2* |
| *AFF3* | *ANKRD33B* | *ARPP19* |
| *ATRN* | *ANKRD34C* | *BTG2* |
| *BRI3BP* | *APOD* | *CNDP1* |
| *BTBD3* | *C4orf50* | *CPQ* |
| *C11orf87* | *CAMK1G* | *CTHRC1* |
| *CACNA1E* | *CARTPT* | *CYB5R1* |
| *CAMK1D* | *CD163* | *EDN3* |
| *CCDC3* | *CIT* | *ENPEP* |
| *CDKL1* | *CLSTN2* | *FZD6* |
| *CLASP1* | *DUSP4* | *HCLS1* |
| *CNKSR3* | *FKBP5* | *HLA-DRA* |
| *CSMD1* | *FOSL2* | *HMGCS1* |
| *CTXN3* | *GRB10* | *IFI44* |
| *CYP39A1* | *IGFBP6* | *LRRC17* |
| *DCLK1* | *IGSF9B* | *MMRN2* |
| *EMP1* | *JRK* | *NXNL2* |
| *ENPP6* | *KDM5A* | *OAS2* |
| *ETV5* | *KLHL3* | *P2RY13* |
| *FAM126B* | *KSR2* | *PI16* |
| *FBXO33* | *LAD1* | *SFRP4* |
| *FOXN3* | *LCN9* | *ST6GAL2* |
| *FRRS1L* | *LRP4* | *TUSC3* |
| *FRY* | *LRRC53* |  |
| *GFOD1* | *LTB* |  |
| *GFRA1* | *MAP3K9* |  |
| *GRAMD1B* | *MAPK4* |  |
| *HECW1* | *MAS1* |  |
| *HERC1* | *MED28* |  |
| *HOMER1* | *MKLN1* |  |
| *HSPA4L* | *MLEC* |  |
| *HTT* | *PAPLN* |  |
| *HYOU1* | *PLCH1* |  |
| *IGFBP5* | *PTK2B* |  |
| *ITPR1* | *RHBDL3* |  |
| *KCNA1* | *SHC3* |  |
| *KCNA3* | *SIK2* |  |
| *KCNB1* | *SLC36A1* |  |
| *KCND3* | *SPEN* |  |
| *KCNH1* | *STRBP* |  |
| *KCNS1* | *SYNJ2* |  |
| *KCNS2* | *TET3* |  |
| *KDM7A* | *TNS1* |  |
| *KIAA1549* | *UNC13A* |  |
| *KIAA1671* | *UNC79* |  |
| *LPL* | *USH1C* |  |
| *LRRC8B* | *VEGFA* |  |
| *LUZP1* | *VPS53* |  |
| *MAP9* | *WIPF3* |  |
| *MGAT5* | *ZBTB16* |  |
| *NAV1* | *ZBTB40* |  |
| *NOMO2* | *ZNF189* |  |
| *NUAK1* |  |  |
| *PCSK1* |  |  |
| *PDE4DIP* |  |  |
| *PDP1* |  |  |
| *PDZD2* |  |  |
| *PEX5L* |  |  |
| *PIK3CA* |  |  |
| *PLEKHM3* |  |  |
| *PLS1* |  |  |
| *PLXNA2* |  |  |
| *PLXNA4* |  |  |
| *PRKCA* |  |  |
| *PROS1* |  |  |
| *PTPRJ* |  |  |
| *RAPGEF5* |  |  |
| *RGS4* |  |  |
| *RORB* |  |  |
| *SCN8A* |  |  |
| *SCRN1* |  |  |
| *SHISA6* |  |  |
| *SLC24A2* |  |  |
| *SLC38A1* |  |  |
| *SLC38A2* |  |  |
| *SORL1* |  |  |
| *SORT1* |  |  |
| *SPTLC3* |  |  |
| *SRC* |  |  |
| *STX1B* |  |  |
| *STXBP5L* |  |  |
| *SV2C* |  |  |
| *TCP11L1* |  |  |
| *TDRD9* |  |  |
| *THRB* |  |  |
| *TMX4* |  |  |
| *TNR* |  |  |
| *TRIO* |  |  |
| *TTBK2* |  |  |
| *UGGT1* |  |  |
| *USP37* |  |  |
| *VPS13D* |  |  |
| *VSIG10L* |  |  |
| *WDFY3* |  |  |
| *WIF1* |  |  |
| *ZDHHC23* |  |  |

**Table S3.** **Sex-stratified main effects of refined Blue expression-based polygenic score (ePGS) and adversity exposure score on depressive (PHQ-9) and anxiety (GAD-7) symptoms in the UK Biobank.**

| **ePGS** | **Outcome** | **Females β (p)** | **Males β (p)** | **N (Females)** | **N (Males)** |
| --- | --- | --- | --- | --- | --- |
| **Blue module ePGS/ACC** | PHQ-9 | 0.010 (0.10) | 0.001 (0.82) | 81,453 | 64,034 |
|  | GAD-7 | 0.010 (0.13) | 0.002 (0.59) | 81,700 | 64,237 |
| **Early adversity** | PHQ-9 | 0.25 (<10^-16^) | 0.25 (<10^-16^) | 80,877 | 63,780 |
|  | GAD-7 | 0.22 (<10^-16^) | 0.21 (<10^-16^) | 81,120 | 63,981 |
| **Adult adversity** | PHQ-9 | 0.23 (<10^-16^) | 0.27 (<10^-16^) | 57,888 | 46,153 |
|  | GAD-7 | 0.20 (<10^-16^) | 0.22 (<10^-16^) | 57,990 | 46,241 |
| **Combined adversity** | PHQ-9 | 0.29 (<10^-16^) | 0.33 (<10^-16^) | 57,312 | 45,899 |
|  | GAD-7 | 0.24 (<10^-16^) | 0.27 (<10^-16^) | 57,410 | 45,985 |

β values denote standardized regression coefficients from linear regression models, with p values shown in parentheses. All models were adjusted for age, genotype array, assessment centre, and 40 genetic principal components.

**Table S4.** **Gene–environment interactions effects between refined Blue module ePGS/ACC and adversity on PHQ-9 and GAD-7 symptoms in the UK Biobank (females).** Related to Figure 4.

| **Outcome** | **Adversity** | **Females β (SE)** | **P-value** | **FDR** | **N (Females)** |
| --- | --- | --- | --- | --- | --- |
| PHQ-9 | Early | 0.011 (0.003) | 0.0004 | 0.002* | 80,877 |
|  | Adult | 0.009 (0.004) | 0.01 | 0.020* | 57,888 |
|  | Combined | 0.011 (0.004) | 0.0015 | 0.005* | 57,312 |
| GAD-7 | Early | 0.008 (0.003) | 0.01 | 0.023* | 81,120 |
|  | Adult | 0.001 (0.004) | 0.75 | 0.75 | 57,990 |
|  | Combined | 0.005 (0.004) | 0.17 | 0.20 | 57,410 |

β values denote standardized regression coefficients for the interaction term; standard errors (SE) are shown in parentheses. Nominal p values and FDR-adjusted p values (Benjamini–Hochberg) are reported. FDR correction was applied within sex across all interaction tests reported for both outcomes (PHQ-9 and GAD-7). All models were adjusted for age, genotype array, assessment centre, and 40 genetic principal components.

**Table S5. Gene–environment interactions effects between refined Blue module ePGS/ACC and adversity on PHQ-9 and GAD-7 symptoms in the UK Biobank (males).**

| **Outcome** | **Adversity** | **Males β (SE)** | **P-value** | **FDR** | **N (Males)** |
| --- | --- | --- | --- | --- | --- |
| PHQ-9 | Early | 0.002 (0.004) | 0.62 | 0.63 | 63,780 |
|  | Adult | 0.007 (0.005) | 0.22 | 0.42 | 46,153 |
|  | Combined | 0.008 (0.005) | 0.13 | 0.42 | 45,899 |
| GAD-7 | Early | 0.002 (0.004) | 0.63 | 0.63 | 63,981 |
|  | Adult | 0.006 (0.005) | 0.28 | 0.42 | 46,241 |
|  | Combined | 0.006 (0.005) | 0.22 | 0.42 | 45,985 |

β values denote standardized regression coefficients for the interaction term; standard errors (SE) are shown in parentheses. Nominal p values and FDR-adjusted p values (Benjamini–Hochberg) are reported. FDR correction was applied within sex across all interaction tests reported for both outcomes (PHQ-9 and GAD-7). All models were adjusted for age, genotype array, assessment centre, and 40 genetic principal components.

**Table S6. Gene–environment interaction effects between Green and Tan module ePGS/ACC and adversity on depressive (PHQ-9) and anxiety (GAD-7) symptoms in UK Biobank females.**

| **ePGS** | **Outcome** | **Adversity** | **Females β (SE)** | **p-value** | **FDR** | **N (Females)** |
| --- | --- | --- | --- | --- | --- | --- |
| **Green module ePGS/ACC** | PHQ-9 | Early | 0.001 (0.003) | 0.87 | 0.998 | 80,877 |
|  |  | Adult | -0.002 (0.004) | 0.53 | 0.998 | 57,888 |
|  |  | Combined | -0.002 (0.004) | 0.62 | 0.998 | 57,312 |
|  | GAD-7 | Early | 0.002 (0.003) | 0.53 | 0.998 | 81,120 |
|  |  | Adult | -0.0002 (0.004) | 0.96 | 0.998 | 57,990 |
|  |  | Combined | 0.00001 (0.004) | 0.10 | 0.998 | 57,410 |
| **Tan module ePGS/ACC** | PHQ-9 | Early | 0.004 (0.003) | 0.16 | 0.32 | 80,877 |
|  |  | Adult | 0.007 (0.004) | 0.05 | 0.32 | 57,888 |
|  |  | Combined | 0.005 (0.004) | 0.16 | 0.32 | 57,312 |
|  | GAD-7 | Early | 0.002 (0.003) | 0.56 | 0.68 | 81,120 |
|  |  | Adult | 0.002 (0.004) | 0.54 | 0.68 | 57,990 |
|  |  | Combined | 0.001 (0.004) | 0.74 | 0.74 | 57,410 |

β values denote standardized regression coefficients for the interaction term, with standard errors (SE) shown in parentheses. Nominal p values and FDR-adjusted p values (Benjamini–Hochberg) are reported. All models were adjusted for age, genotype array, assessment centre, and 40 genetic principal components. FDR correction was applied across all interaction tests, including both outcomes (PHQ-9 and GAD-7).

**Table S7. Item-level interaction effects between refined Blue module ePGS/ACC and adversity on PHQ-9 symptoms in UK Biobank females.** Related to Figure 5.

| **PHQ-9 item** | **Adversity type** | **β (SE)** | **P-value** | **FDR** | **N** |
| --- | --- | --- | --- | --- | --- |
| **Suicidality** | Early-life | 0.0123 (0.003) | 6.7×10⁻⁵ | 0.002* | 81,542 |
|  | Adult | 0.0089 (0.004) | 0.009 | 0.034* | 58,174 |
|  | Combined | 0.0135 (0.004) | 1.9x10^-4^ | 0.003* | 57,570 |
| **Depressed mood** | Early-life | 0.0076 (0.003) | 0.016 | 0.034* | 81,731 |
|  | Adult | 0.0063 (0.004) | 0.094 | 0.144 | 58,276 |
|  | Combined | 0.0091 (0.004) | 0.014 | 0.034* | 57,658 |
| **Concentration problems** | Early-life | 0.0099 (0.003) | 0.001 | 0.008* | 81,862 |
|  | Adult | 0.0093 (0.004) | 0.010 | 0.034* | 58,326 |
|  | Combined | 0.0119 (0.004) | 0.001 | 0.008* | 57,703 |
| **Feelings of inadequacy** | Early-life | 0.0094 (0.003) | 0.003 | 0.013* | 81,666 |
|  | Adult | 0.0062 (0.004) | 0.104 | 0.148 | 58,230 |
|  | Combined | 0.0092 (0.004) | 0.014 | 0.034* | 57,618 |
| **Anhedonia** | Early-life | 0.0047 (0.003) | 0.122 | 0.164 | 81,761 |
|  | Adult | 0.0090 (0.004) | 0.013 | 0.034* | 58,283 |
|  | Combined | 0.0106 (0.004) | 0.003 | 0.013* | 57,666 |
| **Sleep problems** | Early-life | 0.0042 (0.003) | 0.181 | 0.222 | 81,811 |
|  | Adult | 0.0071 (0.004) | 0.057 | 0.103 | 58,312 |
|  | Combined | 0.0062 (0.004) | 0.096 | 0.144 | 57,688 |
| **Tiredness** | Early-life | 0.0078 (0.003) | 0.012 | 0.034* | 81,799 |
|  | Adult | 0.0037 (0.004) | 0.318 | 0.322 | 58,307 |
|  | Combined | 0.0067 (0.004) | 0.065 | 0.110 | 57,684 |
| **Changes in appetite** | Early-life | 00034 (0.003) | 0.303 | 0.322 | 81,830 |
|  | Adult | 0.0047 (0.004) | 0.228 | 0.258 | 58,308 |
|  | Combined | 0.0047 (0.004) | 0.229 | 0.258 | 57,685 |
| **Psychomotor changes** | Early-life | 0.0058 (0.003) | 0.056 | 0.103 | 81,843 |
|  | Adult | 0.0037 (0.004) | 0.322 | 0.322 | 58,302 |
|  | Combined | 0.0052 (0.004) | 0.153 | 0.196 | 57,683 |

All models tested the interaction between refined Blue module ePGS/ACC and adversity on standardized PHQ-9 item scores in UK Biobank females. Models were adjusted for age, genotype array, assessment centre, and 40 genetic principal components. β represents the interaction term coefficient and SE its standard error. Exact p-values are reported. FDR correction (Benjamini–Hochberg) was applied across item-level interaction tests within females. Sample size varies across models due to the availability of adversity measures.

**Table S8. Description of early-life and adult adversity scores derived from UK Biobank questionnaire items.**

| Early-life adversity score | |
| --- | --- |
| Question asked: “*When I was growing up*…” | UK Biobank  data-field |
| 1. I felt loved | 20489, 29076 |
| 2. People in my family hit me so hard that it left me with bruises or marks | 20488, 29077 |
| 3. I felt that someone in my family hated me | 20487, 29078 |
| 4. Someone molested me (sexually) | 20490, 29079 |
| 5. There was someone to take me to the doctor if I needed it | 20491, 29080 |
| Adult adversity score | |
| Question asked: “*Since I was sixteen*…” |  |
| 1. I have been in a confiding relationship | 20522 |
| 2. A partner or ex-partner deliberately hit me or used violence in any other way | 20523 |
| 3. A partner or ex-partner repeatedly belittled me to the extent that I felt worthless | 20521 |
| 4. A partner or ex-partner sexually interfered with me, or forced me to have sex against my wishes | 20524 |
| 5. There was money to pay the rent or mortgage when I needed it | 20225 |

Items were selected from the UK Biobank touchscreen questionnaires. Protective items were reverse coded before score computation. Early-life and adult adversity scores were computed separately by summing standardized item responses. A combined adversity score was derived by summing the standardized early-life and adult adversity scores.
